# A hybrid receptor binding protein enables phage F341 infection of *Campylobacter* by binding to flagella and lipooligosaccharides

**DOI:** 10.1101/2023.09.05.556331

**Authors:** Line Jensen Ostenfeld, Anders Nørgaard Sørensen, Horst Neve, Amira Vitt, Jochen Klumpp, Martine C. Holst Sørensen

## Abstract

Flagellotropic bacteriophages are interesting candidates as therapeutics against pathogenic bacteria dependent on flagellar motility for colonization and causing disease. Yet, phage resistance other than loss of motility has been scarcely studied. Here we developed a soft agar assay to study flagellotropic phage F341 resistance in motile *Campylobacter jejuni*. We found that phage adsorption was prevented by diverse genetic mutations in the lipooligosaccharides forming the secondary receptor of phage F341. Genome sequencing showed phage F341 belongs to the *Fletchervirus* genus otherwise comprising capsular-dependent *C. jejuni* phages. Interestingly, phage F341 encodes a hybrid receptor binding protein (RBP) predicted as a short tail fiber showing partial similarity to RBP1 encoded by capsular-dependent *Fletchervirus,* but with a receptor binding domain similar to tail fiber protein H of *C. jejuni* CJIE1 prophages. Thus, *C. jejuni* prophages may represent a genetic pool from where lytic *Fletchervirus* phages can acquire new traits like recognition of new receptors.

## Introduction

Flagella-driven motility of pathogenic bacteria is considered an important virulence factor as it is essential for establishing gut colonization, as well as mediate epithelial cell adhesion (Haiko and Westerlund-Wikström, 2013). In addition, flagella also function as initial attachment sites for many bacteriophages (phages), the natural predators of bacteria (Evans et al., 2010a; Evans et al., 2010b; Guerrero-Ferreira et al., 2011; Iino and Mitani, 1967; Lotz et al., 1977; Raimondo et al., 1968; Zhilenkov et al., 2006; Pate et al., 1979; Choi et al., 2013; Baldvinsson et al., 2014). It is believed that by binding to flagella, phages increase their capture range and thus increase the chances of finding suitable hosts in complex environments like the gut. Such phages are generally referred to as flagellotropic phages, and they are considered interesting candidates for therapeutic applications against pathogenic bacteria relying on motility for causing disease.

Most phages attaching to flagella require flagellar rotation to establish infection promoting the translocation of the phage along the flagella filament down to the bacterial surface like a “nut-on-a- bolt” (Ravid and Eisenbach, 1983; Schade et al., 1967; Yen et al., 2012; Evans et al., 2010a; Samuel et al., 1999; Choi et al., 2013; Guerrero-Ferreira et al., 2011; Baldvinsson et al., 2014; Sørensen et al., 2015). Several studies propose that when flagellotropic phages reach the bacterial body they interact with a secondary receptor, where injection of the phage genome takes place (Bender et al., 1989; Schade et al., 1967; Lotz et al., 1977; Samuel et al., 1999; Guerrero-Ferreira et al., 2011; Pate et al., 1979; Baldvinsson et al., 2014). Yet, only very few studies describe the nature of the secondary receptor such as pili recognized by *Caulobacter* phage ϕCbK, lipopolysaccharides recognized by *Agrobacterium* phage 7-7-1 and the AcrABZ-TolC complex proposed for *Salmonella* phage Chi (Guerrero-Ferreira et al., 2011; Gonzales et al., 2018; Esteves et al., 2021). Also, only in the case of phage 7-7-1 have receptor binding proteins (RBPs) been proposed for flagellotropic phages (Gonzales and Sharf, 2021). Furthermore, little is known about how resistance may develop under conditions where there is a selective pressure to maintain motility, such as during host colonization. Often *in vitro* selection and characterization of phage resistant mutants is used as a method to identify both phage resistance mechanisms and surface components important for phage infection. Yet, *in vitro* experiments investigating flagellotropic phage resistance development often result in loss of motility or mutations in flagella associated genes (Choi et al., 2013; Evans et al., 2010b; Lotz et al., 1976; Coward et al., 2006). As a result it is difficult to identify secondary surface structures important for flagellotropic phage infection *in vitro* and to study resistance development that may better reflect *in vivo* conditions.

Flagellotropic phages have been proposed as a promising intervention strategy against the major foodborne human pathogen *Campylobacter jejuni* that rely on flagellar motility for establishing intestinal colonization and human disease (Hammerl et al., 2014; El-Shibiny et al., 2009; Carvalho et al., 2010). Most of the flagellotropic *C. jejuni* phages described so far belong to the *Firehammervirus* genus, but no specific surface components nor receptor binding proteins have been identified for these phages. In addition, a study reporting resistance development towards flagellotropic *C. jejuni* phages describes loss of bacterial motility (Coward et al., 2006). Thus, it is currently unknown how resistance may develop towards these phages when there is a selective pressure to maintain motility such as during colonization of the host. *C. jejuni* phage F341 is dependent on motility for successful host infection and binds to the flagella using the distal tail fibers (Baldvinsson et al., 2014). Phage F341 can however also infect a non-motile mutant of its propagation host *C. jejuni* NCTC12658 with a low efficiency supporting the interaction with a secondary receptor present on the bacterial body (Baldvinsson et al., 2014). Yet, the specific surface components recognized by phage F341 as well as its genetic content and genetic relationship with other *C. jejuni* phages have not been determined. Here we aimed to develop an *in vitro* assay to study phage resistance development against flagellotropic phages when motility is maintained using *C. jejuni* and phage F341 as model organisms. We further aimed to (i) identify the secondary receptor of phage F341, (ii) determine its genetic content and (iii) identify the receptor binding proteins responsible for host recognition.

## Materials and Methods

### Bacteria and bacteriophage strains and standard growth conditions

All bacterial strains and information on phage F341 are listed in table S1. *C. jejuni* was standardly grown on base II plates with 5% calf blood (BA) at 37°C under microaerobic conditions (6% O2, 6% CO2, 88% H2N2 or 6% O2, 6% CO2, 88% N2).

The genome sequences of *Campylobacter jejuni* strain NCTC12658 and *Campylobacter* phage F341 are deposited in NCBI Genbank under the NZ_CP081480.1 and OQ864999 Accession numbers. The fastq sequencing files of the LJ variants can be found at European Nucleotide Archive under project Accession number PRJEB61440.

### Soft agar assay

*C. jejuni* strains of interest were grown under standard growth conditions and harvested into cation adjusted (1 mM CaCl2, 10 mM MgSO4) Brain Hearth Infusion broth (Oxoid) (CBHI) and subsequently adjusted to OD600 of 0.01 in 10 ml cultures. Phage F341 was added to reach a multiplicity of infection (MOI) of 0.001 and the cultures were incubated for 24 hours under standard growth conditions. Following incubation, serial dilutions (10^-4^-10^-8^) were made in phosphate buffered saline and 100 μl of each serial dilution was plated onto BA plates with 5 ml of NZCYM overlay agar at 0.6% agar concentration that had been pre-dried for 45 min under a flow hood. The plates were then incubated under standard growth conditions for 2-3 days until single colonies were visible. Swarming colonies were re-streaked on regular BA plates, harvested into BHI and stored at -80 °C. Swarming colonies were later tested for phage sensitivity and motility as mentioned below.

### Motility assay

Motility assays were performed as described previously (Sørensen et al., 2011). In brief, *C. jejuni* strains of interest were grown at standard conditions for 18-24 hours. Cells were then harvested into BHI and adjusted to an OD600 of 0.1. One microliter of the suspension was spotted in the center of five Heart infusion broth (HIB) (Difco) 0.25% agar plates that had been pre-dried for 45 min in a flow hood. Plates were incubated under standard growth conditions and growth zones representing motility were measured after 24, 42 and 48 hours. Measurements and standard deviations represent the mean counts from two independent experiments.

### Phage F341 propagation and titration, standard and antibody plaque assay

Phage F341 was propagated on *C. jejuni* NCTC12658 as described previously using the double-layer agar method (Baldvinsson et al., 2014). Phages from lysed plates were harvested into 5 ml SM buffer (50 mM Tris-HCl, pH-7.5, 100 mM CaCl, 8 mM MgSO4) and stored at 4°C. Titration and standard plaque assays were performed as previously described using spot assays on bacterial lawns (Sørensen et al., 2021a). Briefly, tenfold serial dilutions of the F341 phage stock were made in SM buffer up to 10^−7^, and three aliquots of 10 μl of the undiluted stock (10^0^) and each dilution were spotted on bacterial lawns with the strains of interest. Plates were incubated for 18–24 h at 37°C under standard *C. jejuni* growth conditions and plaques were counted and the mean plaque forming units per ml (pfu/ml) were calculated. The RBP antibody plaque assay was performed as previously described (Sørensen et al., 2021a). Briefly, anti-F341_RBP serum and pre-serum (serum extracted prior to immunization) was added at a 1:10 ratio to the phage F341 stock for 1 hour at room temperature before serial dilutions were made and spotted on NCTC12658 lawns. All experiments were performed in duplicate and the data presented are the means from two independent experiments.

### Phage adsorption assay

Phage adsorption assays were performed as previously described (Baldvinsson et al., 2014). Briefly, *C. jejuni* cultures were washed and adjusted to an OD600 of 0.4 in CBHI and inoculated with phage F341 at an MOI of 0.001. Cultures were incubated at 37°C and samples containing free-phages were collected at 0, 30, 60, and 90 min and filtered through an 0.22-μm sterile syringe filter (Millipore). Filtered samples were stored at 4°C until enumeration (plaque assay). Free phages were enumerated on *C. jejuni* NCTC12658 and calculated as the percentage of free phages compared to time point zero for each sample (phage pfu/ml at time zero was accounted as 100%). The data are representative of two independent experiments and represent the mean percentages of free phages and standard deviations thereof.

### DNA isolation

Genomic DNA from phage F341 was isolated as described previously (Sørensen et al., 2021a). The DNA concentration was estimated visually based on 0.7-1 % agarose gel electrophoresis as *Campylobacter* phages contain modified DNA (Crippen et al., 2019) that cannot be measured correctly by fluorometric quantification. Chromosomal DNA from *C. jejuni* NCTC12658 was isolated using Maxwell 16 Tissue DNA extraction kit (Promega) (Illumina sequencing) and the MagAttract HMW DNA kit (Qiagen) (SMRT sequencing, Pacific Biosciences) according to the manufacturer’s instructions. Chromosomal DNA from the LJ variants was isolated using Maxwell 16 Tissue DNA extraction kit (Promega) according to manufacturer’s instructions.

### Genome sequencing and bioinformatic analyses

Genome sequencing of *C. jejuni* NCTC12658 was done on the Illumina platform (Illumina Hiseq and Miseq) and on a PacBio RS II device (Pacific Biosciences, Menlo Park, CA, USA) using P6/C4 chemistry using one SMRT cell. The complete *de novo* genome was assembled using SMRT Analysis version 2.3 and the HGAP3 algorithm. Average coverage was 634-fold. The genome was annotated using the NCBI Prokaryotic Genome Annotation Pipeline (PGAP). Genome sequencing of phage F341 resistant NCTC12658 variants LJ1, LJ6, LJ8, LJ9, LJ10, LJ11 and LJ13 was performed using the Illumina platform (Illumina Miseq). Sequencing of LJ4 was not performed due to failed library preparation. Basic variant detection analysis was performed using CLC genomics workbench version 21.0.3 using default settings. Reads mapping to the NCTC12658 reference genome were as follows for the phage F341 resistant variants; LJ1: 384.052, LJ6: 674.826, LJ8: 656.167, LJ9: 535.977, LJ10: 565.008, LJ11: 537.104 and LJ13: 460.315 with coverage of detected SNP’s ranging from 10-fold to 132-fold. *De novo* assembly of the phage F341 resistant variants was performed using CLC genomic workbench version 21.0.3 with a total number of contigs generated: 144 (LJ1), 157 (LJ6), 141 (LJ8), 173 (LJ9), 113 (LJ10), 151 (LJ11) and 39 (LJ13).

The phage F341 genome was sequenced on a PacBio RS II device (Pacific Biosciences, Menlo Park, CA, USA) using P6/C4 chemistry using one SMRT cell. Sequencing produced 30591 reads with a mean read length of 25853. The complete *de novo* genome was assembled using SMRT Analysis version 2.3 and the HGAP3 algorithm. Average coverage of the phage F341 genome was 6006-fold. The phage genome was annotated using CPT Galaxy and WebApollo platforms (Afgan et al., 2018). Comparative genomics of *Fletchervirus* phages was performed in CLC main workbench version 20.0.4 using the whole genome alignment 20.1 plugin tool with default settings. Upper comparison gradient: ANI (average nucleotide identity), lower comparison: AP (alignment percentage). Detailed *in silico* analyses of phage F341 proteins was performed using InterPro (Mitchell et al., 2019), BLASTp (Johnson et al., 2008), HHpred (Zimmermann et al., 2018), Colab AlphaFold2 (Mirdita et al., 2022) and Dali server (Holm, 2022). Proteins are visualized using Pymol (Schrodinger, 2015).

### Pulsed Field Gel Electrophoresis (PFGE)

PFGE analyses of *C. jejuni* NCTC12658 and LJ variants was performed according to Ribot et al., 2001 using two restriction enzymes, SmaI and KpnI (New England Biolabs), separately.

### Construction of NCTC12658Δ*05515-05525* (NCTC12658ΔLOS) LOS deletion mutant

The deletion of *05515-05525* in NCTC12658 was constructed by replacing most of genes (the first 287 bp of *05525* and the last 36 bp of *05515* are maintained) with a kanamycin (kan) cassette by homologous recombination. A vector was constructed by fusing three PCR amplicons as described below using In-fusion cloning (Takara) and used for homologous recombination. Two PCR fragment (746 bp and 829 bp) flanking the region upstream and downstream the *05515-05525* deletion was amplified from *C. jejuni* NCTC12658 using the primers LOS_F_up and LOS_R_up and LOS_F_down and LOS_R_down, respectively. The kan gene (1398 bp) was amplified from pBCα3 (Bijlsma et al., 1999) using the primers Kan+LOS_F and Kan+LOS_R. The kan gene was then inserted between the upstream and downstream *05515-05525* flanking PCR fragments and simultaneously inserted into the pGEM-7Zf vector (Promega) in the BamHI and XhoI sites using the In-Fusion kit according to the manufacturer’s instructions. The resulting plasmid pMP810 was transformed into *E. coli* StellarTM Competent Cells (Takara) according to manufacturer’s instructions and verified by sequencing. Purified pMP802 plasmid was electroporated into *C. jejuni* NCTC12658 promoting homologous recombination of the flanking *05515-05525* regions to create NCTC126581′*05515-05525*. The mutant was verified by PCR using the LOS_control_F and LOS_control_R primers and sequencing. Plasmids and primers are listed in Table S1.

### Construction of the pMP35 synthetic *F341_rbp* expression plasmid

A 1.439 bp fragment including *F341_rbp* and the downstream *F341_034 chaperone* plus an additional TAA stop codon was synthetically produced and inserted into the NdeI and XhoI cloning sites of pET28a+ producing the F341_RBP protein with an N-terminal his-tag used for purification of the protein.

### F341_RBP protein expression, purification and antibody production

The F341_RBP was expressed and purified as previously described using *E. coli* BL21-CodonPlus (DE3)-RIL competent cells (Agilent Technologies) and 0.5 mM IPTG (isopropyl β-D-1- thiogalactopyranoside) induction (Sørensen et al, 2021a). The F341_RBP was purified using Ni^2+^- NTA-affinity chromatography (His GraviTrap, GE Healthcare) according to manufacturer’s instructions and exchanged into PBS buffer using 10 kDA Amicon® Ultra-15 Centrifugal filter units. The purified protein was used to raise polyclonal rabbit antibodies generated by Davids Biotechnologie GmbH (Germany). Anti-F341_RBP serum was purified on a protein A column and stored at -20°C until use. The binding ability of F341_RBP to phage F341 was confirmed by standard Western blotting.

### Transmission electron microscopy and immunogold labeling

Immunogold electron microscopy was performed as described previously (Sørensen et al, 2021a). Staining was performed as previously reported with 2% (w/v) uranyl acetate on freshly prepared carbon films (Sørensen et al., 2015). Grids were analyzed in a Tecnai 10 transmission electron microscope (TEM) (FEI Thermo Fisher, Eindhoven, The Netherlands) at an acceleration voltage of 80 kV. Micrographs were taken with a MegaView G2 charge-coupled device camera (EMSIS, Muenster, Germany).

## Results

### A soft agar assay allows *in vitro* isolation of motile phage resistant mutants of *C. jejuni*

To investigate phage resistance in *C. jejuni* against flagellotropic phages when motility is maintained, we developed a soft agar assay that would allow us to screen for motile phage resistant variants *in vitro*. We exposed *C. jejuni* NCTC12658 to phage F341 in liquid culture for 24 hours followed by plating serial dilutions on double layered soft agar (SA) plates where swarming/motility of single colonies could be observed (Figure 1A). To evaluate motility levels, we also plated the *C. jejuni* NCTC12658 wildtype (motile) and a NCTC12658Δ*motA* mutant (non-motile) as controls. As expected, many colonies of the NCTC12658 wildtype exhibited swarming behavior, while no swarming was observed for the NCTC12658Δ*motA* colonies (Figure 1A). Most colonies originating from NCTC12658 exposed to phage F341 demonstrated no swarming behavior, consistent with previous findings that loss of motility rapidly occurs when *C. jejuni* is exposed to flagellotropic phages *in vitro*. Still, a number of motile colonies could be observed (Figure 1A), suggesting that phage F341 resistance without loss of motility can also develop *in vitro*. To verify that the swarming colonies exposed to phage F341 indeed were fully motile and resistant to phage F341, we isolated 14 swarming colonies from the SA plate (named LJ1 to LJ14) and investigated phage F341 sensitivity by plaque formation and assessed motility in soft agar. Six of the LJ colonies were still highly motile but had not gained phage resistance, as F341 plaque formation was comparable to the NCTC12658 wildtype (data not shown). However, eight colonies (LJ1, LJ4, LJ6, LJ8, LJ9, LJ10, LJ11 and LJ13) demonstrated complete resistance to phage F341 as no lysis nor plaque formation was observed, while still exhibiting motility at levels comparable to the NCTC12658 wildtype (Figure 1B). To determine if phage F341 resistance was associated with lack of phage binding, we investigated whether F341 could adsorb to the eight motile phage resistant LJ strains (Figure 1C). Our data showed that for all eight LJ strains, phage adsorption was prevented suggesting changes in surface structures such as alteration or masking of the receptor recognized by phage F341. Thus, our soft agar assay allowed us to isolate motile *C. jejuni* NCTC12658 variants *in vitro* exhibiting complete resistance towards flagellotropic phage F341.

**Figure 1.**
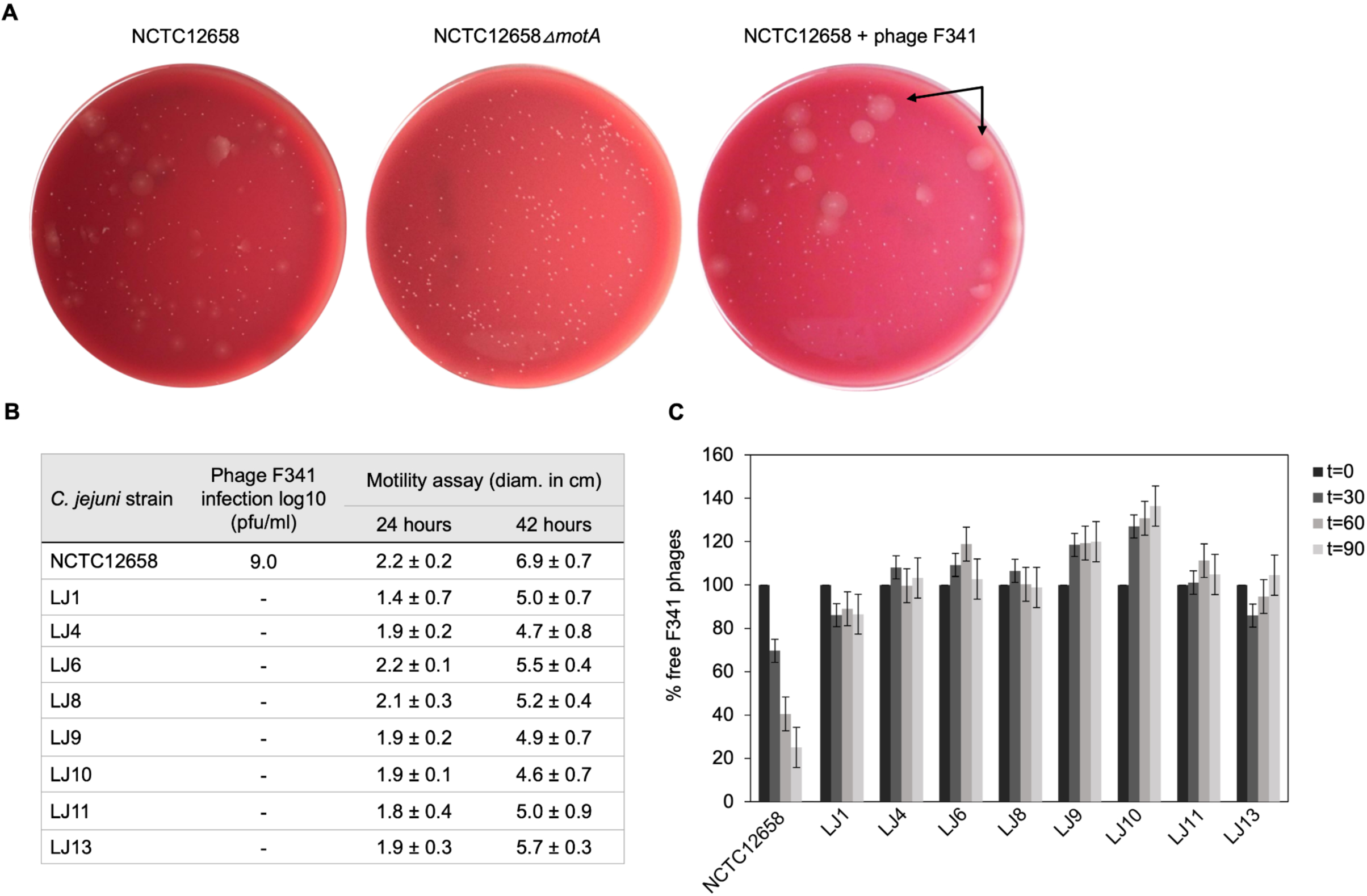
Motile phage F341 resistant *C. jejuni* NCTC12658 variants (LJ strains) isolated *in vitro* demonstrate lack of phage binding. (A) Soft agar (SA) assay with *C. jejuni* NCTC12658, NCTC12568Δ*motA* and NCTC12658 exposed to phage F341 demonstrating different levels of swarming and non-swarming colonies. Examples of swarming colonies spreading out on the agar plate by flagellar motility are indicated by arrows (B) Phage F341 infection and motility profiles of isolated swarming NCTC12658 colonies exposed to phage F341 (LJ strains). (C) Phage F341 adsorption assay of phage resistant LJ strains. t equals the timepoint in minutes where samples were collected, and the number of free phages determined. Pfu/ml of free phages at time zero was accounted as 100%. The images represent two independent experiments; however, the LJ colonies were only isolated from plates from one of the independent experiments. Error bars and ± indicated mean standard deviations.

### Phage F341 resistant NCTC12658 variants contain only few genetic changes mostly associated with the lipooligosaccharide (LOS) locus

To identify genetic changes that could account for phage F341 resistance, we sequenced the genomes of seven phage resistant NCTC12658 variants (LJ1, LJ6, LJ8, LJ9, LJ10, LJ11 and LJ13) with Illumina sequencing. We also genome sequenced the NCTC12658 wildtype strain using both Illumina and Pacific Bioscience (PacBio) SMRT sequencing to fully assemble the genome. We then used the complete genome of NCTC12658 and the sequencing reads of the seven phage resistant variants to detect single nucleotide polymorphisms (SNPs) and larger gene insertions or deletions (Table 1, Table S2 and S3). To identify potential genomic rearrangements we also performed *de novo* assembly and pulsed field gel electrophoresis (PFGE), but no rearrangements were observed (Figure S1).

**Table 1.** Single nucleotide polymorphisms (SNPs) and gaps observed in motile phage F341 resistant NCTC12658 variants in the lipooligosaccharide (LOS) locus following basic variant detection analysis. See also Table S2 and S3 and Figure S1.

| *C. jejuni*<br>strain | SNP or<br>gap<br>position | Gene/s affected | Product (homolog in *C. jejuni*<br>NCTC11168) | Proposed function | SNP description<sup>a,b)</sup> |
| --- | --- | --- | --- | --- | --- |
| LJ1 | 1.073.110 | *K5A08_05515* | Glycosyltransferase (Cj1135,<br>two-domain β-1,4<br>glucosyltransferase) | Attaches β-1,4 glucose residues<br>to heptose in the inner LOS core | A deletion (100%), early stop:<br>280 aa vs 515 aa (SND) |
| LJ6 | 1.073.464 | *K5A08_05515* | Glycosyltransferase (Cj1135,<br>two-domain β-1,4<br>glucosyltransferase) | Attaches β-1,4 glucose residues<br>to heptose in the inner LOS core | 8 A's (100%) – gene off<br>(polyA tract variation) |
| LJ8 | 1.079.283 | *cstIII*<br>(*K5A08_05540*) | α-2,3-sialyltransferase (CstIII, α-<br>2,3 sialyltransferase) | Attaches N-acetylneuraminic<br>acid (sialic acid) to galactose in<br>the outer LOS core | 7 A's (91%), truncation: 261 aa<br>vs 294 aa<br>(polyA tract variation) |
| LJ11 | 1.075.897 | *K5A08_05525* | Capsular polysaccharide<br>synthesis protein (Cj1137, α-<br>1,2-galactosyltransferase) | Attaches α-1,2 galactose<br>residues to galactose in the outer<br>LOS core | 7 A's (100%) – gene off<br>(PolyA tract variation) |
| LJ10 | 1.074.429..<br>1.076.390 | *K5A08_05520*<br>*K5A08_05525* | Glycosyltransferase family 2<br>protein (Cj1136, β-1,3-galactosyltransferase)<br>Capsular polysaccharide<br>synthesis protein (Cj1137, α-<br>1,2-galactosyltransferase) | Cj1136: Attaches β-1,3<br>galactose (outer core) to heptose<br>(inner core)<br>Cj1137: Attaches α-1,2<br>galactose residues to galactose<br>in the outer LOS core | Most of *K5A08_05520* and all<br>of *K5A08_05525* have no read<br>coverage (gene deletions) |
| LJ13 | 1.074.820..<br>1.076.414 | *K5A08_05520*<br>*K5A08_05525* | Glycosyltransferase family 2<br>protein (Cj1136, β-1,3-galactosyltransferase)<br>Capsular polysaccharide<br>synthesis protein (Cj1137, α-<br>1,2-galactosyltransferase) | Cj1136: Attaches β-1,3<br>galactose (outer core) to heptose<br>(inner core)<br>Cj1137: Attaches α-1,2<br>galactose residues to galactose<br>in the outer LOS core | Most of *K5A08_05520* and all<br>of *K5A08_05525* have no read<br>coverage (gene deletions)<sup>c)</sup> |

<sup>a)</sup> SNV: single nucleotide variation, SND: single nucleotide deletion. The frequency of the SNPs observed are provided in percentages.
<sup>b)</sup> On and off indicate the gene expression status as a result of polymeric tract changes causing frame shift mutations.
<sup>c)</sup> *De novo* assembly confirms that genes *K5A08\_05520* and *K5A08\_05525* are no longer present in the genome of the LJ13 variant (data not shown).

As the genome of LJ9 was not fully covered by the sequencing we could not identify genetic changes associated with phage F341 resistance in this variant. However, our analysis identified a limited number of SNPs (≤14) and two gene deletions in the remaining phage resistant NCTC12658 variants (Supplementary Results, Table 1, Table S2 and S3). Most SNPs were associated with insertions or deletions (indels) in homopolymeric G (polyG) or A (polyA) tracts causing phase variable gene expression in *C. jejuni*. Such indels are expected within a given *C. jejuni* population due to the stochastic nature of slipped strand mispairing of such regions during DNA replication (Bayliss et al., 2012; van der Woude, 2011). Thus, some SNPs may not be due to phage F341 exposure but could be a random mutation event originating from picking single LJ colonies. Also, different polymeric tract lengths were in some cases observed between the two sequencing methods for the NCTC12658 wildtype. Hence, when gene expression was found to be both turned on and off in the wildtype NCTC12658 strain sensitive to phage F341, or when different polymeric tract lengths where observed that did not lead to differences in the gene expression state, the SNP was not considered relevant. In addition, our experimental data showed that phage resistance was associated with lack of phage adsorption, indicating genetic changes associated with surface structures. Genetic changes not considered relevant for phage F341 resistance development are presented in Supplementary Results and Table S2 and S3.

*C. jejuni* is highly glycosylated on its surface and produce capsular polysaccharides (CPS), lipooligosaccharides (LOS), N-linked glycosylated outer membrane proteins and O-linked glycosylated flagellin forming the flagella filament (Burnham and Hendrixson, 2018). Notably, four out of the seven LJ strains (LJ1, LJ6, LJ8 and LJ11) contain SNPs in genes encoding sugar transferases modifying the LOS structure of *C. jejuni* NCTC12658 (Table 1). NCTC12658 contains the same LOS locus as *C. jejuni* NCTC11168 producing an inner LOS core containing ketodeoxytonic acid, heptose, phoshoethanolamine and glucose, and an outer LOS core containing N-acetylgalactoseamine, N- acetylneuraminic acid (sialic acid), glucose and galactose (Figure 2A) (Karlyshev et al., 2005). LJ6, LJ8 and LJ11 all contain polyA tract variations that either switch off gene expression or lead to the formation of truncated protein products of *K5A08_05515, ctsIII* and *K5A08_05525,* respectively. These genes are responsible for modifying the outer LOS core with galactose and sialic acid, and the inner LOS core with glucose residues suggesting that these LJ strains contain diverse LOS structures compared to the NCTC12658 wildtype (Table 1, Figure 2A). As polymeric tracts found at the gene end in *C. jejuni* have been proposed to influence transcription and/or translation of the downstream gene (Kim et al., 2012), the polyA tract variation of *cstIII* in LJ8 may instead affect the downstream *neuB1* involved in sialic acid biosynthesis (Table 1). A single nucleotide deletion was observed also in the *K5A08_05515* gene in LJ1 introducing an early stop and a truncated protein product. Furthermore, large gaps were found in the LOS loci covering the *K5A08_05520* and *K5A08_05525* gene region in LJ10 and LJ13 (Table 1, Table S3). The deletion of these two genes in the LOS locus was furthermore confirmed by the *de novo* assembly of LJ13 (data not shown). *K5A08_05520* has been proposed to link the outer LOS core to the inner LOS core. Thus, six out of the seven phage F341 resistant NCTC12658 variants show diverse genetic changes that may all result in a different LOS structure as compared to the wildtype NCTC12658 (Figure 2A).

**Figure 2.**
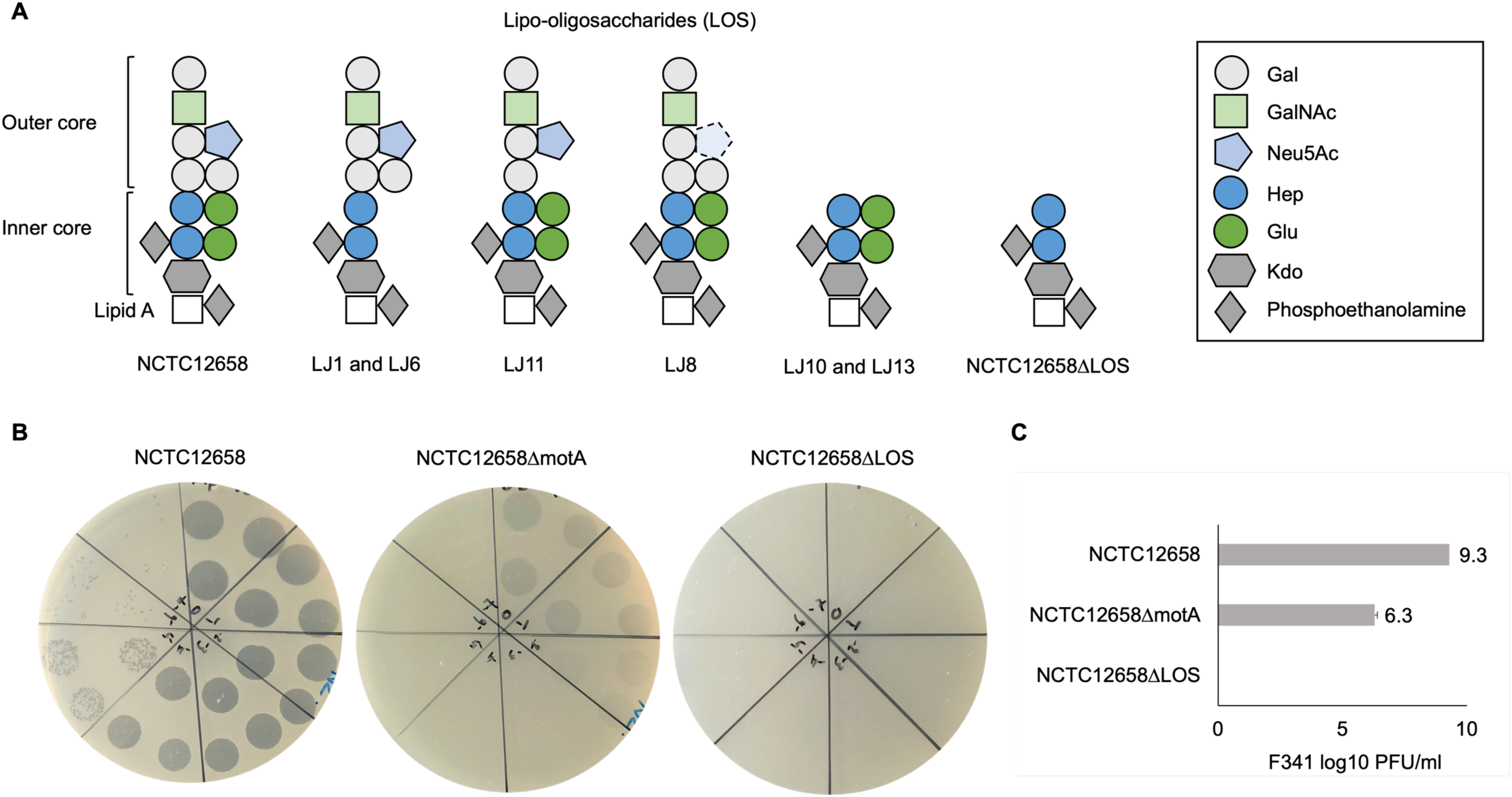
Phage F341 uses lipooligosaccharides as a secondary receptor for successful infection of its host *C. jejuni* NCTC12658. (A) Schematic representation of the lipooligosaccharides (LOS) putatively produced by *C. jejuni* NCTC12658, the LJ phage F341 resistant NCTC12658 variants and the *C. jejuni* NCTC126581′LOS (*C. jejuni* NCTC126581′*05515-05525*) mutant. (B) Standard plaque assays of phage F341 on its propagation host *C. jejuni* NCTC12658, the non-motile *C. jejuni* NCTC126581′*motA* mutant and *C. jejuni* NCTC126581′LOS. (C) Mean F341 log10 pfu/ml values from standard plaque assays on *C. jejuni* NCTC12658, *C. jejuni* NCTC126581′*motA* and *C. jejuni* NCTC126581′LOS. 3 x 10 μl of undiluted phage F341 stock and tenfold serial dilutions up to 10^-7^ are spotted clockwise from the top of all plaque assay plates. The images represent 2-4 independent experiments and error bars indicated standard deviations. Gal: galactose, GalNAc: N-acetylgalactoseamine, Neu5Ac: N-acetylneuraminic acid (sialic acid), Hep: heptose, Glu: glucose, Kdo: ketodeoxytonic acid. See also Figure S2.

### Phage F341 is dependent on LOS for infection of *C. jejuni* NCTC12658

As the genome analysis of the phage F341 resistant motile NCTC12658 variants showed several genetic changes in the LOS locus, we speculated if phage F341 could be dependent on LOS as a secondary receptor for successful host infection. Many of the SNPs and gene deletions were found in the *K5A08_05515 - K5A08_05525* gene region responsible for both modifying and linking the inner and the outer LOS core. We therefore created a *C. jejuni* NCTC126581′*05515-05525* (NCTC126581′LOS) deletion mutant only producing the inner LOS core lacking modifying glucose residues (Figure 2A). Before conducting further experiments, we confirmed that the NCTC126581′LOS mutant was still motile at levels similar to the NCTC12658 wildtype (Figure S2). We then investigated phage F341 sensitivity by plaque formation and compared the results to those obtained on the NCTC12658 wildtype and the non-motile NCTC126581′*motA* mutant. While phage F341 infected the non-motile NCTC126581′*motA* mutant with a lower efficiency of plating as demonstrated previously (Baldvinsson et al, 2014), no lysis nor plaque formation was observed on the NCTC126581′LOS mutant (Figure 2B and C). These results demonstrate that phage F341 is completely dependent on the LOS for successful infection thereby functioning as a secondary receptor in *C. jejuni* NCTC12658

### Phage F341 belongs to the *Fletchervirus* genus and demonstrates high sequence similarity to other phages in this genus

To determine the genetic content of *Campylobacter* phage F341, we performed whole genome sequencing. Sequencing revealed a genome size of 131.68 kb with an average GC content of 26%, and BLASTn searches showed that phage F341 belong to the *Fletchervirus* genus. This is consistent with a genome size of approximately 130 kb being characteristic for phages belonging to this genus (Javed et al., 2014). In addition, phage F341 encodes five tRNA’s and a large number (12) of putative homing endonucleases which has also previously been observed in other *Fletchervirus* phages (Javed et al., 2014). To further investigate the genetic relationship between phage F341 and other *Fletchervirus* phages, we performed comparative genomics using the genome sequences of 18 *Fletchervirus* phages in our collection (Sørensen et al., 2021a). Comparative genomics showed 89-96% average nucleotide identity (alignment percentage: 85-94%, average: 88,4%) between the F341 genome and the 18 *Fletchervirus* genome sequences (Figure 3A) with the closest relative being phage F336 (alignment percentage: 92%, average 96%). Thus, phage F341 shows high sequence similarity to other *Fletchervirus* phages and we were only able to detect five genes (*F341_034*, *F341_035*, *F341_109*, *F341_128* and *F341_135*) that were either only observed in phage F341, or which showed no or very low sequence similarity to genes/translated proteins encoded by other *Fletchervirus* phages (Table 2 and Supplementary Table S4). Further detailed *in silico* analyses demonstrated that F341_109 and F341_128 show structural similarity to baseplate wedge proteins. F341_109 also demonstrates structural similarity to adaptor proteins forming base plate attachment sites for tail fibers in other phages (Supplementary Table S4). *F341_135* is likely encoding fibritin which in *E. coli* phage T4 forms neck/collar fibers and exhibit chaperone function by promoting the assembly of the long tail fibers and their attachment to tail base plate (Tao et al., 1997). Interestingly, this gene was not observed at the corresponding position in other *Fletchervirus* genomes currently available. The remaining two unique F341 genes (*F341_034* and *F341_035*) were located within the receptor binding protein (RBP) region for capsular-dependent *Fletchervirus* phages and will be discussed in more detail in the following paragraph. Thus, only few genes diverge in phage F341 compared to other *Fletchervirus* phages, yet proposed functions suggest differences in tail associated genes involved in host recognition.

**Figure 3.**
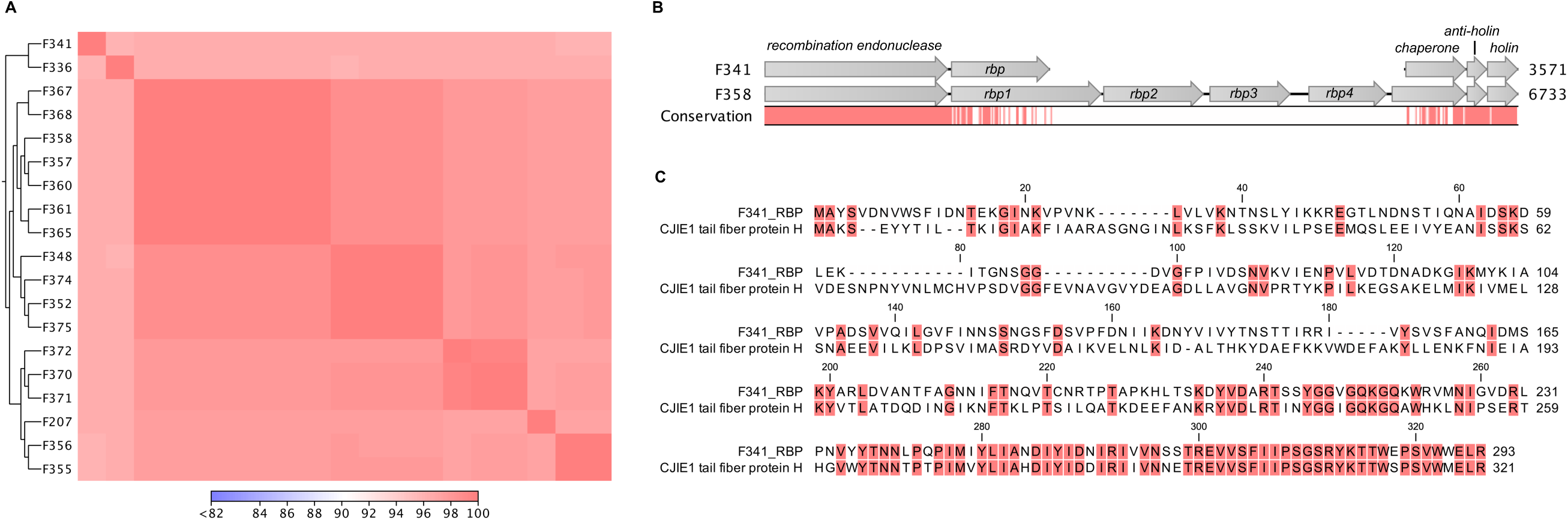
Flagellotropic phage F341 belongs to the *Fletchervirus* genus but shows limited sequence similarity to related phages in the RBP region. (A) Comparative genomics of phage F341 and 18 *Fletchervirus* phages. Whole genome alignment visualized as a heat map for average nucleotide identity. Closest relatives are further indicated by the trees. (B) Alignment of the receptor binding protein region in *Fletchervirus* phages dependent on the capsular polysaccharide for infection and phage F341. Phage F358 is used as a representative of capsular-dependent *Fletchervirus* phages. (C) Protein alignment of the RBP of phage F341 and the tail fiber protein H of the CJIE1 prophage in *C. jejuni* strain HF5-4A-4. Comparative genomics was performed using CLC main workbench version 20.0.4 using the whole genome alignment 20.1 tool with default settings. See also Figure S3 and S4.

**Table 2.**
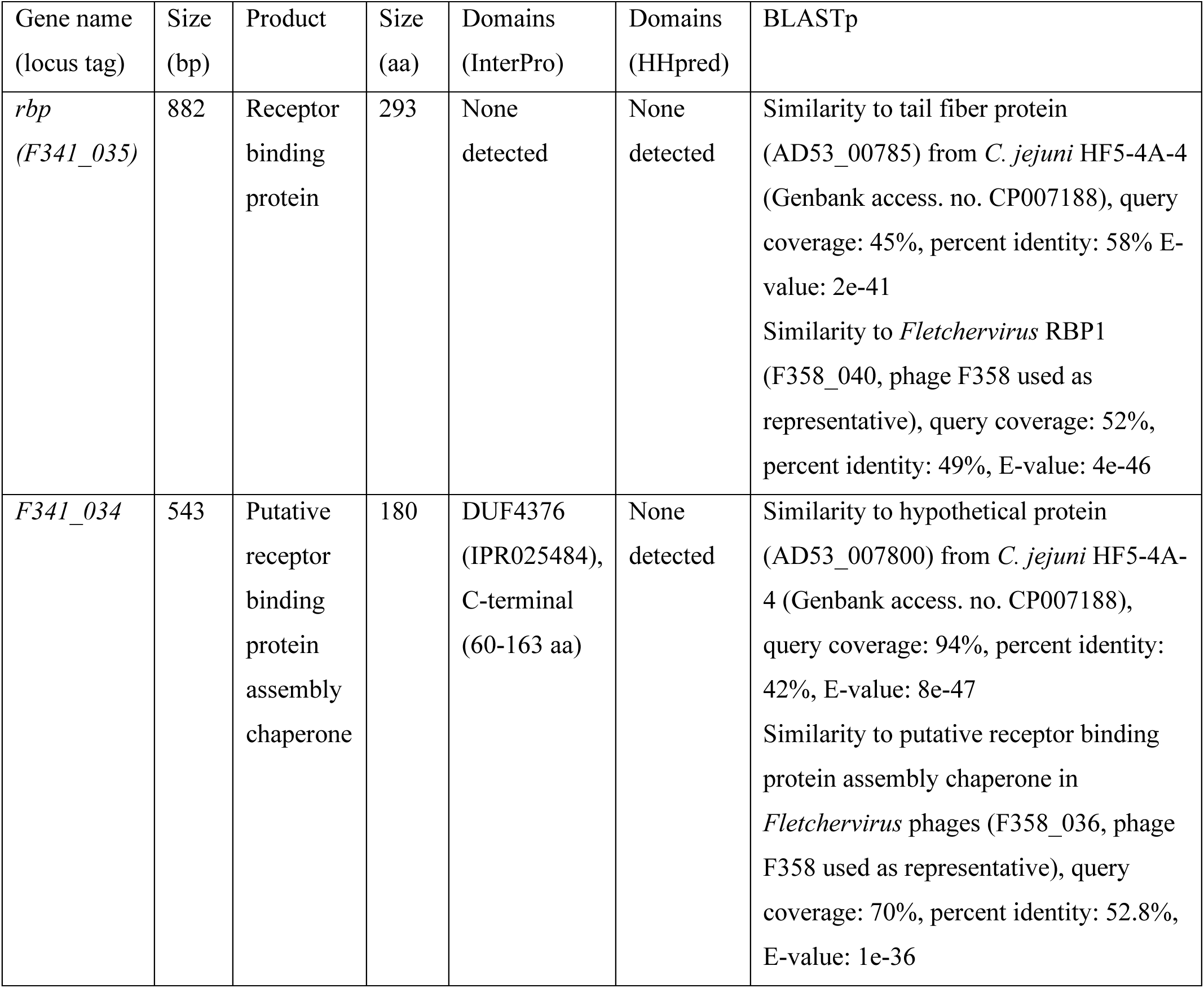
*In silico* analysis of the genes encoded by phage F341 in the *Fletchervirus* RBP region.

### Phage F341 encodes a hybrid receptor binding protein with sequence similarity to both *Fletchervirus* and CJIE1 prophages

*Fletchervirus* phages are usually dependent on capsular polysaccharides for infection of *C. jejuni* and encode up to four receptor binding proteins (*rbp1*-*rpb4*) where RBP1 is responsible for interacting with the common *O*-methyl phosphoramidate phage receptor (Sørensen et al, 2021a). In these phages, the RBPs are located between a conserved putative recombination endonuclease and putative antiholin (Sørensen et al., 2021a). However, phage F341 encodes a unique 882 bp gene (*F341_035*, *rbp*) downstream of the recombination endonuclease, demonstrating no hits to other genes when using BLASTn. When aligning the putative *rbp* gene in F341 to the *rbp* region of capsular-dependent *Fletchervirus* phages, a 26,5% sequence identity to *rbp1* (1.338 bp) could however be observed (Figure 3B). Further protein alignment showed that F341_RBP shares some sequence similarity with the N-terminal of RBP1 up to the point of the pectin lyase domain found in these proteins (Figure S3). Interestingly, BLASTp demonstrated sequence similarity to the C-terminal of the tail fiber protein H encoded by the cryptic CJIE1 prophage in *C. jejuni* strain HF5-4A-4 (Figure 3C, Table 2). Complete alignment of the protein sequences of F341_RBP and the tail fiber protein H showed an almost identical sequence in the far C-terminal of these two RBPs (Figure 3C). These results suggest that F341 encodes a novel hybrid RBP with similarity to RBP1 encoded by the lytic capsular-dependent *Fletchervirus* phages and the RBP (tail fiber protein H) of a CJIE1 prophage.

In both the *Fletchervirus* phages and the CJIE1 prophages, a putative chaperone for correct RBP folding is found downstream of the RBPs containing a DUF4376 domain in the C-terminal (Sørensen et al., 2021a; Zampara et al., 2021). Also downstream of the putative *rbp* gene in phage F341, a hypothetical protein encoded by gene (*F341_034*) with a DUF4376 domain in the C-terminal was observed (Table 2). The *F341_034* gene shows sequence similarity in the N-terminal region to a putative chaperone located next to of the tail fiber protein H in *C. jejuni* HF5-4A-4, whereas the C-terminal region shows similarity to the putative chaperone located downstream of the *rbp* genes in capsular-dependent *Fletchervirus* phages (Table 2, Figure S4). Thus, these results suggest that *F341_034* encodes a putative novel hybrid chaperone associated with the correct folding of the F341 RBP.

### The hybrid receptor binding protein of phage F341 is essential for phage infection and is structurally predicted as a short tail fiber

To experimentally prove the function of the proposed RBP encoded by phage F341, we raised polyclonal antibodies against the purified protein and used these in an antibody plaque assay to test inhibition of phage infection. Also, labeled antibodies where applied during transmission electron microscopy to determine the location of the putative RBP on the phage particle. From these experiments, we could confirm that F341_035 encodes the RBP of phage F341, as phage F341 infection of its host was completely blocked in the presence of the RBP antibodies i.e. no lysis nor plaque formation was observed (Figure 4A and B). Antibodies previously raised against RBP1 from capsular-dependent *Fletchervirus* phage F358 showed no effect on phage F341 infection as expected (Figure 4B). The lack of lysis and plaque formation is similar to what we observed on the NCTC12658 LOS deletion mutant suggesting that the F341 RBP is recognizing and binding to the LOS (Figure 2B). Also, transmission electron microscopy of phage F341 using immunogold labeled RBP antibodies demonstrated binding to the distal tail fibers/spikes (Figure 4C).

**Figure 4.**
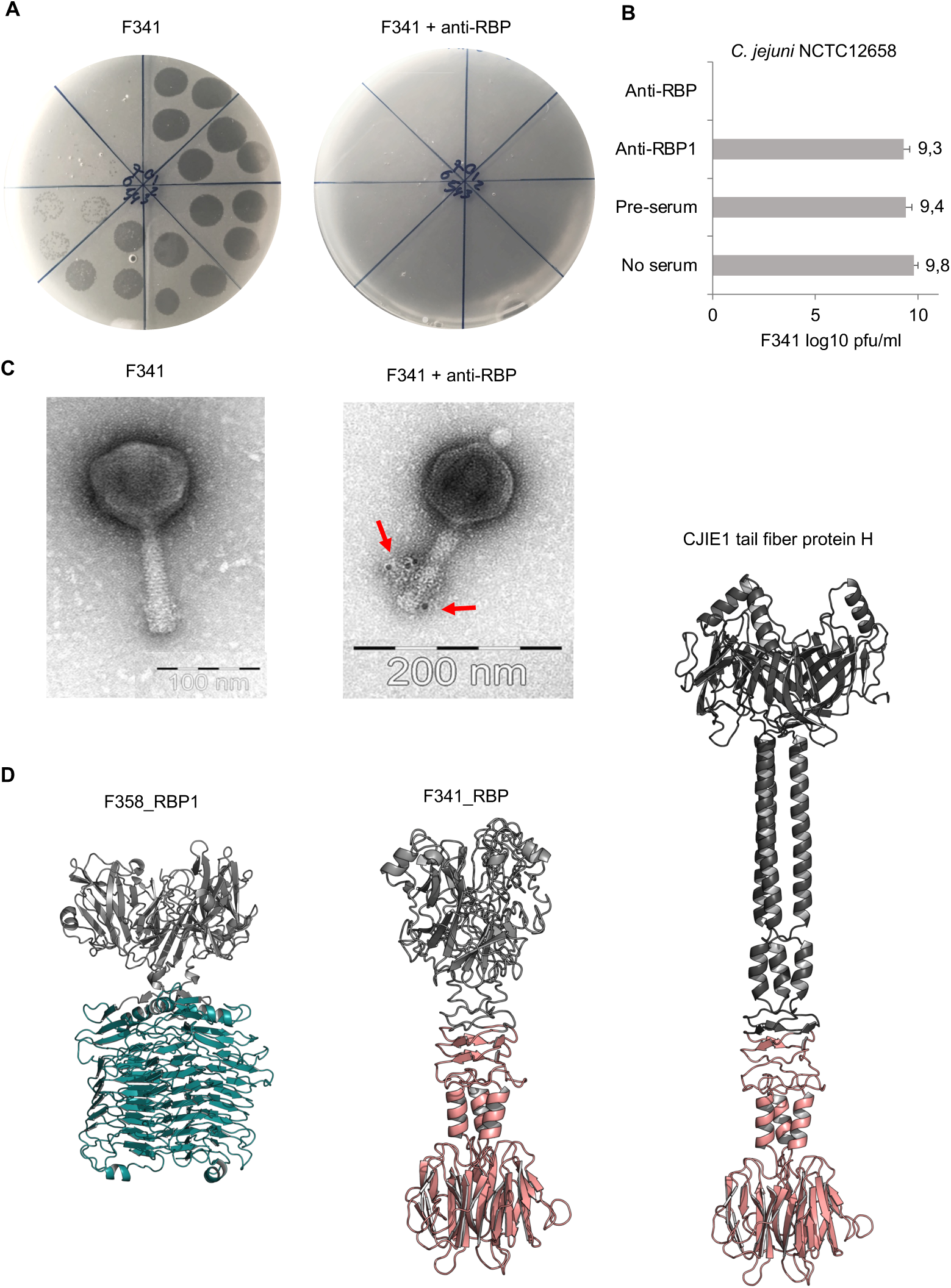
Phage F341 encodes and expresses a unique hybrid RBP with a short tail fiber morphology. (A) Standard and anti-RBP serum plaque assays of phage F341 on its host *C. jejuni* NCTC12658. (B) Mean F341 log10 pfu/ml values from standard and anti-RBP serum plaque assays on *C. jejuni* NCTC12658. Anti-RBP1 is serum produced against RBP1 from capsular-dependent *Fletchervirus* phage F358 (C) Transmission electron microscopy of negatively stained phage F341 and when mixed with immunogold labeled anti-RBP serum. Labelled antibodies bound to phage F341 tail fibers are indicated by arrows. (D) AlphaFold2 predictions of homotrimers of RBP1 from capsular-dependent *Fletchervirus* phage F358, the RBP from flagellotropic *Fletchervirus* phage F341 and the tail fiber protein H of the CJIE1 prophage in *C. jejuni* strain HF5-4A-4. The pectin lyase domain in RBP1 is indicated in cyan blue, whereas the C-terminal region demonstrating sequence similarity between F341_RBP and the tail fiber protein H is indicated in light pink. 3 x 10 μl of undiluted phage F341 stock and tenfold serial dilutions up to 10^-7^ are spotted clockwise from the top of all plaque assay plates. Anti-RBP and anti-RBP1 serum was added at a 1:10 ratio to the phage F341 stock for 1 hour at room temperature before the serial dilutions were prepared and spotted on the NCTC12658 lawn. The images represent 2-4 independent experiments and error bars indicated standard deviations. See also Figure S5.

To further characterize the hybrid RBP of phage F341, we modelled the protein structure using AlphaFold2 (Mirdita et al., 2022). For comparisons, we also modelled the structures of RBP1 from capsular-dependent *Fletchervirus* phage F358 and the tail fiber protein H found in the CJIE1 prophage in the *C. jejuni* HF5-4A-4 genome (Figure 4D). As RBPs often fold into homotrimer, we predicted all three proteins both as monomers and homotrimers (Figure 4D and S5). The F358 RBP1 homotrimer demonstrates a typical tailspike protein (TSP) morphology i.e. short, rigid and stocky, consisting of two distinct regions, an N-terminal head-like domain and a C-terminal comprised of the pectin lysase domain forming parallel ß-helices. A Dali search (Holm, 2022) comparing protein structures in 3D, further demonstrate high structural similarity between the RBP1 pectin lyase domain and the enzymatic domain of the capsule-degrading tail spike protein (TSP) Gp38 from *Klebsiella* phage KP32 (Squeglia et al., 2020). The CJIE1 tail fiber protein H from *C. jejuni* HF5-4A-4 demonstrates typical tail fiber (TF) morphology i.e. long and slim, with an N-terminal globular head domain, a shaft/stem formed primarily of α-helices and a distal C-terminal knob-like structure consisting of a ß-sandwich. The phage F341 RBP also demonstrates TF morphology similar to the CJIE1 tail fiber protein H although the fiber is much shorter. Overall, the F341 RBP structure consists of an N-terminal head domain followed by short shaft/stem region and a distal C-terminal ß-sandwich knob-like structure. The distal C-terminal demonstrating sequence similarity between F341_RBP and the CJIE1 tail fiber protein H is responsible for forming the ß-sandwich knob-like structure and a short α-helical hinge region connected to the shaft/stem. Thus, these results suggest that the distal C-terminal knob-like region is responsible for the interaction with the surface receptor. To further investigate, if the predicted F341 RBP structure resembles other known structural characterized RBPs, we ran a Dali search (Holm, 2022), but no significant hits where observed (data not shown). However, we would propose that the F341 RBP shows some structural similarity to the short tail fibers encoded by *gp12* in *E. coli* phage T4 and to the tail fibers encoded by *E. coli* phage T7 (van Raaij et al., 2001; Garcia-Doval and van Raaij, 2012). In conclusion, our results show that even though phage F341 belongs to the *Fletchervirus* genus it encodes an unique hybrid RBP with a short tail fiber morphology essential for infection of its *C. jejuni* host.

## Discussion

Flagellotropic phages have been proposed as a promising intervention strategy against pathogenic bacteria that rely on flagellar motility for establishing gut colonization and human disease (Esteves and Scharf, 2022). Yet, little is known about how resistance may develop towards such phages when there is a selective pressure to maintain motility, such as during host colonization, as such conditions are difficult to simulate and study *in vitro*. Also, only few studies describe other surface components in addition to flagella that are important for flagellotropic phage infection (Guerrero-Ferreira et al., 2011; Gonzales et al., 2018; Esteves et al., 2021). Thus, new *in vitro* approaches to study flagellotropic phage-host interactions when motility is maintained are required to evaluate the potential of these microorganisms as therapeutics. Here we developed a soft agar assay that allowed us to isolate motile *C. jejuni* NCTC12658 variants *in vitro* exhibiting complete resistance towards flagellotropic phage F341. By characterizing these variants, we found that phage resistance was associated with mutations in the lipooligosaccharides (LOS) forming the secondary receptor of phage F341. Further genomic characterization of phage F341 showed that it belonged to the *Fletchervirus* genus, normally comprising phages dependent on capsular polysaccharides (CPS) for infection (Sørensen et al., 2021a; Sørensen et al., 2015). However, phage F341 contains a hybrid receptor binding protein (RBP) of both lytic *Fletchervirus* phage and CJIE1 prophage origin promoting the recognition of novel surface receptors for phages of this genus.

We previously found that resistance development in *C. jejuni* against CPS-dependent *Fletchervirus* phages was strictly associated with phase variation of surface structures. This was caused by polyG tract variation in the transferases producing the common MeO*P*N receptor and other CPS modifications both *in vitro* and *in vivo* (Sørensen et al., 2011, Sørensen et al., 2012, Aidley et al., 2017, Gencay et al., 2018). Here we found that resistance development towards *Fletchervirus* phage F341 was also associated with changes in surface structures, although by very diverse genetic changes i.e. gene deletions, point mutations and polyA tract variations although all present in the LOS locus. Several surface structures in *C. jejuni* are prone to phase variation due to the presence of polyG tracts within the reading frame of associated genes often encoding sugar transferases (Parkhill et al., 2000). However, only two genes (*wlaN* and *K5A08_05565*) in the LOS locus of *C. jejuni* NCTC12658 contain such tracts and are phase variably regulated (Linton et al., 2000; Gilbert et al., 2002). *wlaN* encodes a ß-1,3-galactosyltransferase responsible for adding a terminal ß-linked galactose to the outer LOS core, while *K5A08_05565* is a homologue of *cj1144-45* in *C. jejuni* NCTC11168 encoding a putative α-1,4- galactosyltransferase associated with terminal α-linked galactose units (Semchenko et al., 2012). As our results demonstrate that phage F341 most likely is binding to several LOS moieties present both in the outer and inner LOS core, turning off the expression of genes that modify the very distal part of the outer LOS core such as *wlaN* and *K5A08_05565*, may not be sufficient to generate phage F341 resistance. On the contrary, numerous polyA tracts of ≥ 7 A’s are found in many LOS genes and may hence also be prone to phase variation, although a high mutational frequency has not been reported for these. Our results show that phase variation as a results of polyA tract variation is also associated with rapid phage resistance development in *C. jejuni in vitro*. Despite the different genetic mutations, we still only observed phage resistance development in *C. jejuni* against *Fletchervirus* phages linked to surface structure diversity that prevent phage adsorption. One might speculate that this could be a bias from the experimental setup and the *in vitro* laboratory settings. Yet previous work performed in our group using phage F341 and *C. jejuni* NCTC11168 in a chicken model, also demonstrated changes in the LOS structure of *C. jejuni* recovered from the chickens following phage exposure, although the data were not statistically significant (Baldvinsson, unpublished). This illustrates that our simple *in vitro* soft agar assay may still be a valuable tool for screening potential phage resistance development against flagellotropic phages simulating *in vivo* conditions where motility is maintained.

We previously demonstrated that phage F341 interacts with both the flagella and the bacterial body using the distal tail fibers (Baldvisson et al., 2014). Here we found that the distal tail fibers are formed by a hybrid RBP that is structurally predicted as a short tail fiber and essential for the interaction with the secondary LOS receptor. Flagellotropic phage 7-7-1 also use the distal tail fibers for the interaction with both the flagella and the body of its *Agrobacterium* hosts (Lotz et al., 1977). Phage 7-7-1 likewise contains short tail fibers and uses the lipopolysaccharides (LPS) as a secondary receptor (Gonzales et al., 2018). As the flagella in both *C. jejuni* and *Agrobacterium* are highly glycosylated (Zebian et al., 2016; Deakin et al., 1999) it is tempting to speculate if the short tail fibers of phage F341 and 7-7-1 are binding loosely to sugar moieties present on the flagella that may appear similar in structure to those of the secondary LOS and LPS receptors, respectively. Indeed, phage adsorption to flagella filaments is generally reversible supporting a loose interaction (Raimondo et al., 1968, Lotz et al., 1977, Baldvinsson et al., 2014). Although the glycans present on the flagella of *C. jejuni* NCTC12658 have not been determined, the strain encodes an almost identical O-linked flagella glycosylation locus as strain NCTC11168. Previous work has shown that flagellins in *C. jejuni* NCTC11168 are modified with legionaminic acid that indeed is an analog of sialic acid present in the LOS outer core (Zebian et al., 2016; Schoenhofen et al., 2017). Our results suggest that phage F341 interacts with sialic acid in addition to several other types of carbohydrate moieties present in both the inner and outer LOS core, as mutations in associated LOS genes lead to phage resistance. We furthermore propose that the short tail fibers formed by the hybrid receptor binding protein of phage F341 show some structural similarity to short tail fibers encoded by *gp12* in *E. coli* phage T4. The short Gp12 tail fibers of phage T4 bind very firmly and irreversible to the LPS core region of *E. coli* to ensure a fixed structure during DNA injection (Thomassen et al., 2003). Thus, we speculate if multiple LOS binding sites may serve a similar purpose for the hybrid RBP of phage F341. This may in addition allow phage F341 to distinguish between LOS and flagella interaction, thereby preventing premature DNA injection upon interaction with the flagella. Yet, as flagella and flagellar motility is needed for efficient phage F341 infection, the flagella interaction may also be a prerequisite for efficient binding to the secondary receptor e.g. by inducing conformational changes in the tail baseplate. Further studies are needed to show if the hybrid RBP of phage F341 indeed is able to interact with legionaminic acid on the flagella. Yet, our study shed new light on how host recognition and interaction may be accomplished by the distal fibers of flagellotropic phages.

The *Fletchervirus* phages sequenced so far are all highly conserved on the sequence level and only distantly related to other phage genera. Indeed, although phage F341 is dependent on flagella and LOS for host infection compared to the majority of *Fletchervirus* phages that rely on capsular polysaccharides, we could only identify five genes/proteins unique to this phage including the hybrid RBP and putative RBP chaperone. The remaining three genes all appear to take part in the correct folding and attachment of the hybrid RBP to the F341 tail base plate. Interestingly, we found that the hybrid receptor binding protein identified in phage F341 demonstrated similarity to the tail fiber H protein encoded by a CJIE1 prophage. Only few prophages have been observed in *C. jejuni* genomes originally identified as integrated elements CJIE1, CJIE2, and CJIE4 in strain RM1221, where CJIE1 shows a Mu-like structure (Parker et al., 2006). It has not been possible to induce or propagate CJIE1 phages except in rare cases suggesting that the majority of these prophages are no longer functional and rather represent phage remnants thus being cryptic prophages (Clark and Ng, 2008; Tanoeiro et al., 2022). We previously confirmed that the tail fiber protein H functions as a receptor binding protein by using this gene from *C. jejuni* strain CAMSA2147 to engineer pyocins targeting *Campylobacter* (Zampara et al, 2021). We did however not identify the receptor recognized by this protein, although our data showed that it was not linked to capsular polysaccharides nor flagella as seen for the lytic phages infecting *C. jejuni* (Zampara et al., 2021). Comparison of CJIE1 regions in different *C. jejuni* strains show various genetic configurations with an overall high sequence similarity except in the tail fiber protein H encoded gene suggesting that these proteins may interact with diverse receptors (Clark and Ng, 2008). This is contrary to the four RBPs identified in the lytic CPS-dependent *Fletchervirus* phages that so far all show high sequence similarity across different phages (Sørensen et al., 2021a). Many *C. jejuni* strains including NCTC12568 do not contain any prophage elements and previous work suggest that the rate of loss and gain of CJIE1 is quite high showing no apparent correlation with colonization levels or human disease (Barton et al., 2007). Yet, the presence of these prophages in very diverse *C. jejuni* strains suggests an early evolutionary origin as well as a benefit of maintaining prophage associated genes although their role remain indecipherable. It is therefore unclear how phage F341 may have acquired the hybrid receptor binding protein and what environmental pressures may have selected for this unique phage. We previously also observed other genes that are shared among *C. jejuni* prophages and lytic phages although only annotated as hypothetical proteins (Sørensen et al, 2021b). This suggests that acquisition or exchange of genetic content between these groups of phages may be a more common phenomenon. This would indeed allow the otherwise highly conserved *Fletchervirus* phages to acquire novel traits such as those observed for phage F341. However, as only a limited number of lytic *Campylobacter* phages have been sequenced so far, more genomes are needed in order to understand the evolutionary relationship between *C. jejuni* prophages and lytic phages.

In conclusion, we here present a simple *in vitro* assay for investigation flagellotropic phage-host interactions when bacterial motility is maintained simulating *in vivo* conditions. In addition, our work further expands on the current knowledge of flagellotropic phages, but also on our understanding of how *Fletchervirus* phages infecting *C. jejuni* have evolved to accomplish new types of host recognition.

## Supporting information

Supplementary data file set

## Acknowledgements

We are thankful to laboratory technician Vi Phuong Thi Nguyen for general assistance on the experimental work and to Stephen J. Ahern for initial annotation of the phage F341 genome. We thank Dr. Anna Bratus-Neuenschwander and the staff of the Functional Genomics Center Zurich for their excellent technical support on the genome sequencing of phage F341. Angela Back (Max Rubner-Institut) is acknowledged for her technical assistance with the TEM analysis. Finally we thank Professor Lone Brøndsted for valued discussion and input on the manuscript.

The work presented here was supported by the Danish Council of Independent Research (grant 4184- 00109B), The Danish AgriFish Agency of Ministry of Environment and Food (grant 34009-14-0873), and Intralytix, Inc. for financial support. We also acknowledge funding of J.K. by Martin J. Loessner, ETH Zurich.

## Author contributions

Conceptualization, M.C.H.S.; methodology, L.J., H.N., A.V. and M.C.H.S.; investigation, L.J., A.N.S.,

H.N., A.V., J.K. and M.C.H.S., formal analysis, L.J. and M.C.H.S.; writing – original draft, M.C.H.S.; funding acquisition, M.C.H.S.; resources, M.C.H.S., H.N., and J.K.; supervision, M.C.H.S.

