## Supplementary data file set for "A hybrid receptor binding protein enables phage F341 infection of *Campylobacter* by binding to flagella and lipooligosaccharides"

**Supplementary Results: Genetic changes observed in phage F341 resistant NCTC12658 variants not related to phage resistance development**

Genetic changes in the LJ variants not associated with phage F341 resistance are presented in Table S1 and described below. Related to Table 1.

PolyG variation (11 and 12 G's) was observed for many of the LJ variants in gene *K5A08_00240* annotated as a bacteriohemerythrin leading to gene expression being predominantly turned off compared to being turned on (10 G's) in the reference genome sequence of NCTC12658. However, we also observed that this gene was predominantly turned off (9 G's) in the NCTC12658 wildtype when looking only at the sequencing reads from PacBio sequencing. Therefore, this SNP was not considered relevant for phage F341 resistance development.

Also, SNPs were observed in the polyG tract in the Cj0031 type II restriction-modification (RM) system (gene *K5A08_00175*) previously described as an internal phage resistance mechanism in *C. jejuni* when switched on (Anjum et al., 2016). Here, expression of the Cj0031 RM system is turned on in the NCTC12658 wildtype sensitive to phage F341, whereas the SNPs observed in *K5A08_00175* indicate that Cj0031 is predominantly switched off in the phage F341 resistant LJ variants (Table S1). We therefore do not expect this SNP to play a role in generating phage resistance against F341. Also, the lack of phage F341 binding to the LJ variants suggest changes in surface structures as the main cause of phage resistance development and not internal resistance mechanism such as RM systems.

Also, as phage F341 infection is completely independent of the presence or absence of capsule polysaccharides (Baldvinsson et al., 2014) and the SNP observed in the CPS locus (*K5A08_06960*) in LJ13 is thus not relevant for phage F341 resistance.

Some SNPs are found in all or most phage resistant variants like a T to C single nucleotide variation (SNV) in gene *K5A08_01925* encoding a DUF1882 domain-containing protein causing an F to S amino acid change. Further bioinformatic analysis suggest that this protein has enzymatic activity and contains a domain that binds dGTP. However, the SNV mutation is not associated with any predicted active sites nor does is lead to conformational changes of the protein (data not shown). Also, a T to C SNV is observed in gene *K5A08_07545* encoding a methyl-accepting chemotaxis protein, but this SNV does not lead to any changes in the protein sequence and may therefore be a randomly accumulated mutation.

Surprisingly, no SNP mutations or genomic deletions were observed in genes associated with the flagella except for the LJ8 variant (Table S1). Here a SNP in the polyG tract found within gene

*K5A08_06310* encoding a Cj1295 homolog producing the di-O-methylglyceroyl modification of Pse in *C. jejuni* NCTC11168 was found in 82% of the sequencing reads for LJ8 turning off the expression of this gene. However, deletion of this gene in strain NCTC11168 was previously shown to maintain phage F341 sensitivity (Baldvinsson et al., 2014).

LJ9 did not show any apparent SNPs, but several smaller gaps were identified for this variant by our basic variant detection analysis suggesting that several areas of the genome were not fully covered by the sequencing. This was also confirmed by the *de novo* assembly resulting in a larger number of contigs (173) compared to that observed for the other phage F341 resistant variants (see materials and methods). It is therefore not clear what may or may not represent real genetic changes occurring in the LJ9 variant as a result of developing phage F341 resistance.

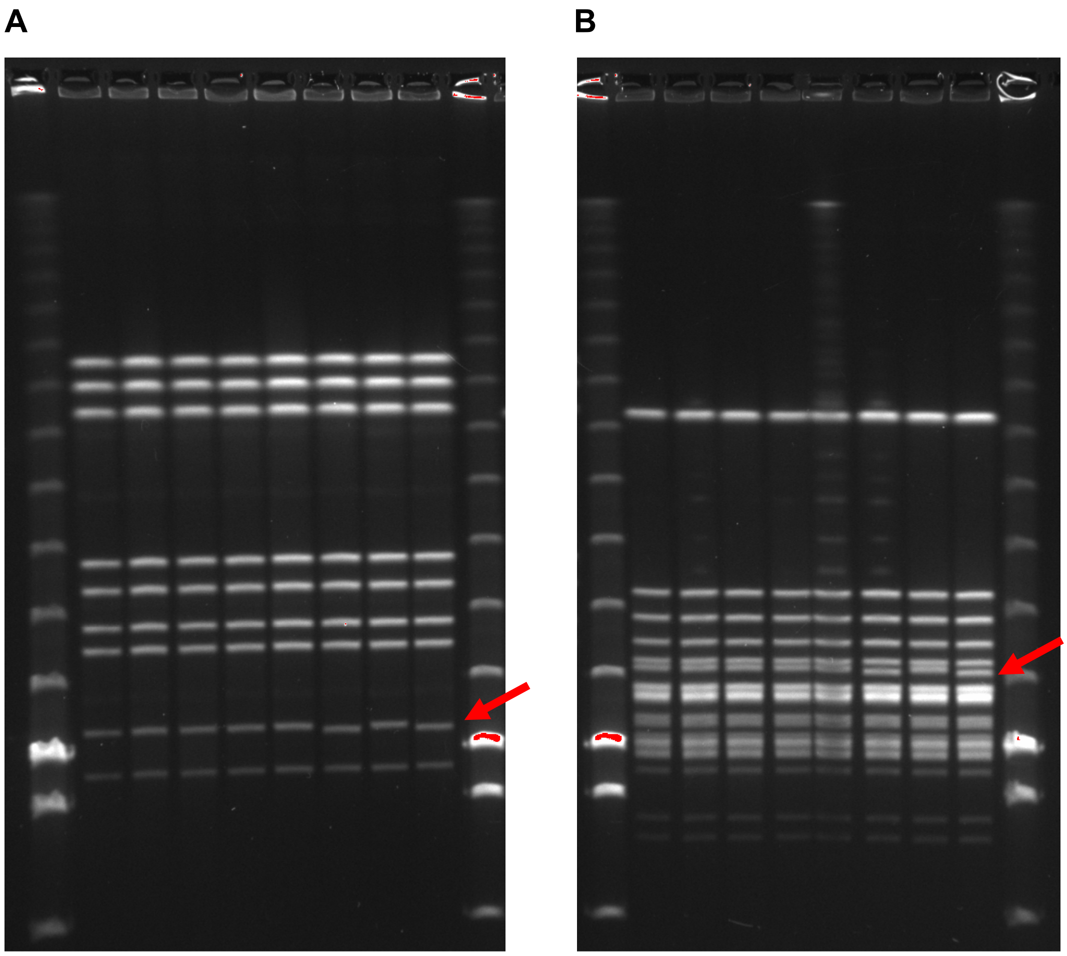


**Figure S1: PFGE analyses of *C. jejuni* NCTC12658 and phage F341 resistant LJ variants.** Related to Table 1 and S2. (A) PFGE of genomic DNA digested with SmaI. From the left lane 1: Lambda marker Medinova NO350S low range, lane 2: NCTC12658, lane 3: LJ1, lane 4: LJ6; lane 5: LJ8, lane 6: LJ9, lane 7: LJ10, lane 8: LJ11, lane 9: LJ13 and lane 10: Lambda marker Medinova NO350S low range. (B) PFGE of genomic DNA digested with KpnI. From the left lane 1: Lambda marker Medinova NO350S low range, lane 2: NCTC12658, lane 3: LJ1, lane 4: LJ6; lane 5: LJ8, lane 6: LJ9, lane 7: LJ10, lane 8: LJ11, lane 9: LJ13 and lane 10: Lambda marker Medinova NO350S low range. Red arrows indicate bands of a slightly smaller size in LJ10 and LJ13 consistent with the deletion of genes *K5A08_05520* and *K5A08_05525* in these strains.

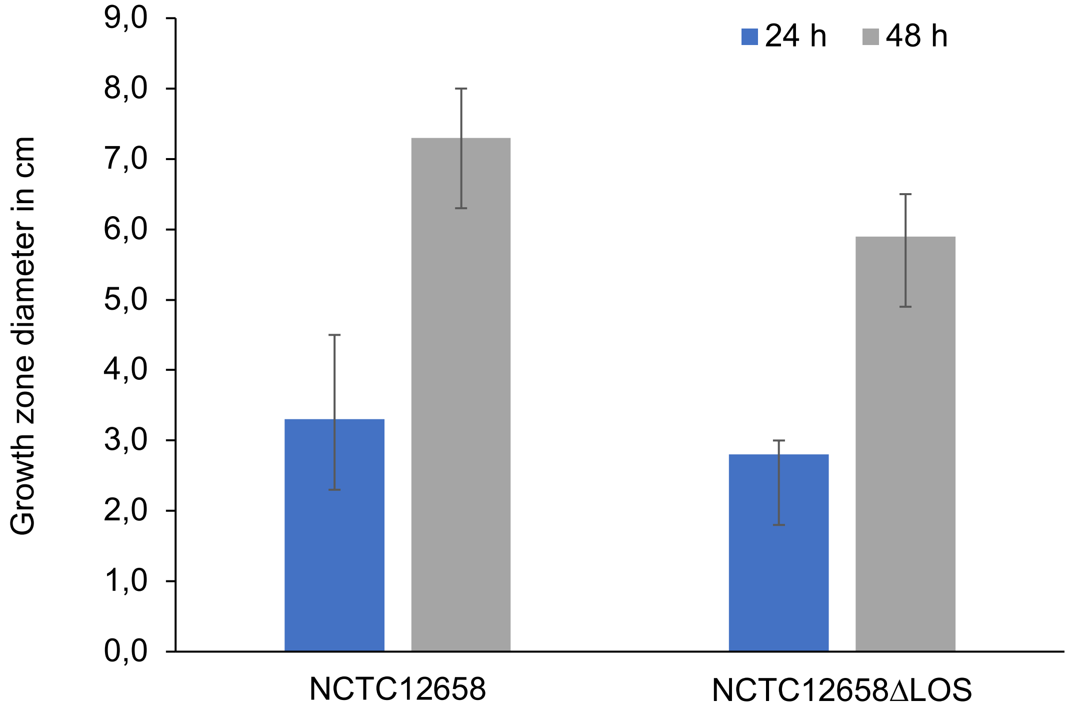


**Figure S2: Motility assay of *C. jejuni* NCTC12658 and NCT12658ΔLOS.** Related to Figure 2. The diameter of growth zones after 24 and 48 hours (h) are indicated. Measurements and standard deviations represent the mean counts from two independent experiments.

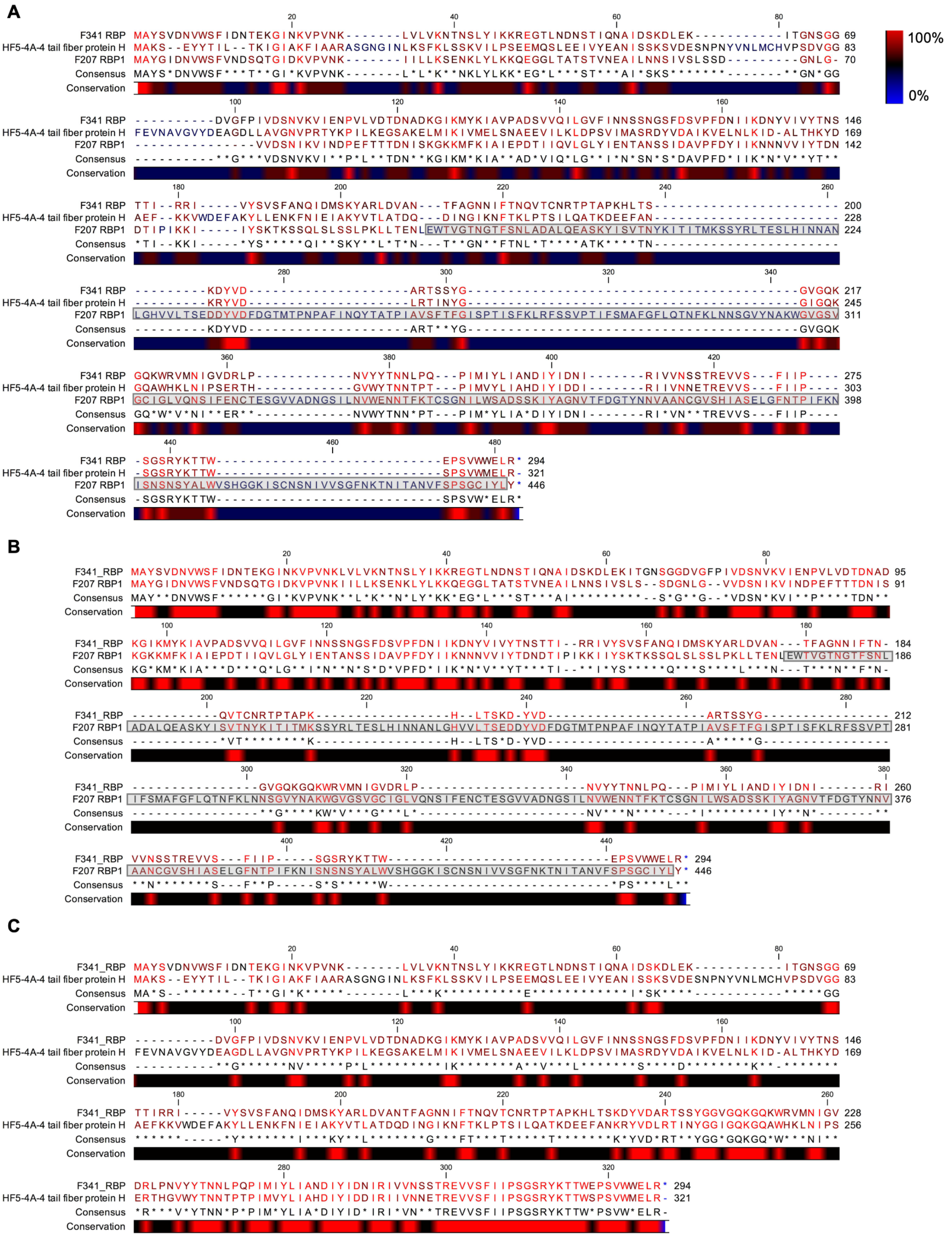


**Figure S3. Protein alignments of *Fletchervirus* RBP1, the CJIE1 tail fiber protein H and the hybrid RBP encoded by phage F341**. Related to Figure 3. (A) Alignment of the RBP from phage F341, RBP1 from capsular-dependent *Fletchervirus* phage F207 and the tail fiber protein H from the CJIE1 prophage found in *C. jejuni* strain HF5-4A-4. The pectin lyase fold found in RBP1 is highlighted in grey. (B) Alignment of RBP from phage F341 and RBP1 from capsular-dependent *Fletchervirus* phage F207. (C) Alignment of RBP from phage F341 and the tail fiber protein H from the CJIE1 prophage found in *C. jejuni* strain HF5-4A-4. Alignment was performed using CLC Main Workbench 20.0.4.

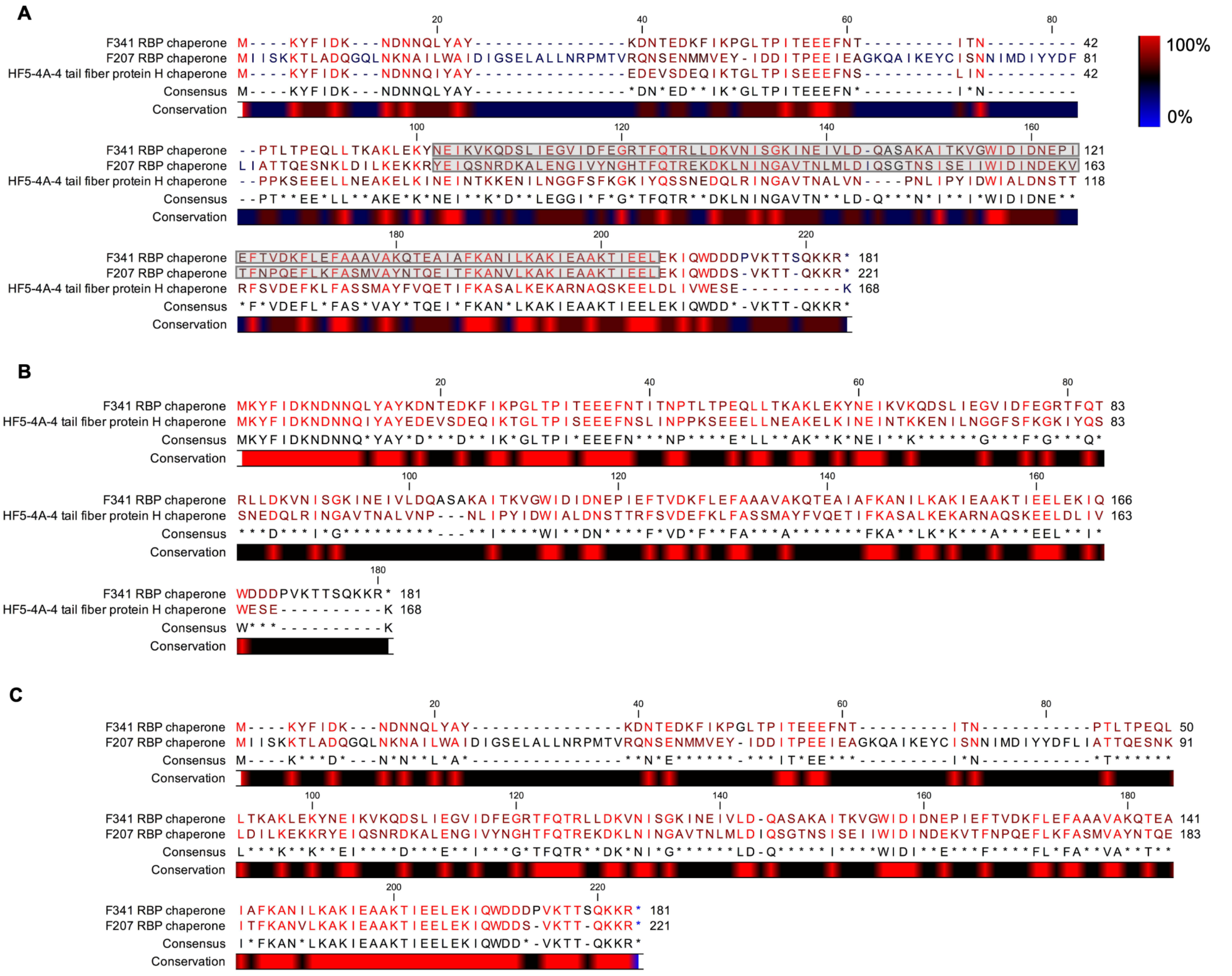


**Figure S4. Protein alignments of the RBP chaperone in capsular-dependent *Fletchervirus* phages, the CJIE1 tail fiber protein H chaperone and the RBP chaperone encoded by phage F341**. Related to Figure 3. (A) Alignment of the RBP chaperone from phage F341, the RBP chaperone from capsular-dependent *Fletchervirus* phage F207 and the tail fiber protein H chaperone from the CJIE1 prophage found in *C. jejuni* strain HF5-4A-4. The DUF4376 (IPR025484) domain observed in the C-terminal of *Fletchervirus* RBP chaperones is highlighted in grey. (B) Alignment of RBP chaperone from phage F341 and the tail fiber protein H chaperone from the CJIE1 prophage found in *C. jejuni* strain HF5-4A-4. C. Alignment of RBP chaperone from phage F341 and RBP chaperone from capsular-dependent *Fletchervirus* phage F207. Alignment was performed using CLC Main Workbench 20.0.4.

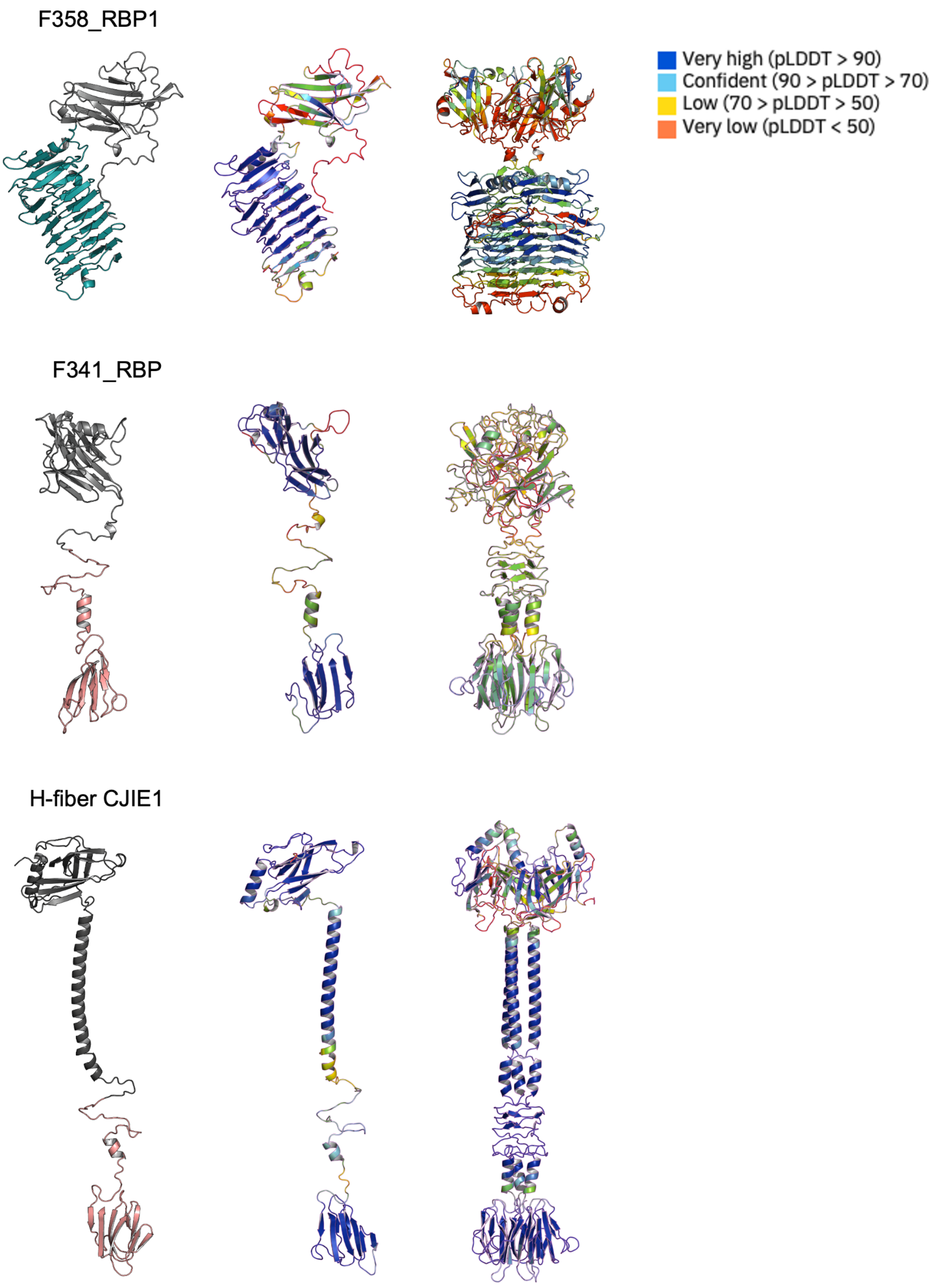


**Figure S5. AlphaFold2 predictions of *Fletchervirus* RBP1, F341_RBP and the tail fiber protein H from the CJIE1 prophage present in *C. jejuni* HF5-4A-4.** Related to Figure 4. AlphaFold2 predictions of monomers and homotrimers of RBP1 from capsular-dependent *Fletchervirus* phage F358, the RBP from flagellotropic *Fletchervirus* phage F341 and the tail fiber protein H of the CJIE1 prophage in *C. jejuni* strain HF5-4A-4. The monomers are colored as presented in the manuscript in Figure 4, i.e. the pectin lyase domain in RBP1 is indicated in cyan blue, whereas the C-terminal region demonstrating sequence similarity between F341_RBP and the tail fiber protein H is indicated in light pink. Both monomers and homotrimers of all three proteins are also shown with the original colors according to the prediction scores as indicated in figure. Proteins are visualized using Pymol.

**Table S1: Key resource table**

| REAGENT or RESOURCE | SOURCE | IDENTIFIER |
|---|---|---|
| Antibodies | | |
| Rabbit polyclonal anti-RBP | This paper | N/A |
| Rabbit polyclonal anti-RBP1 | Sørensen et al., 2021 | N/A |
| Bacterial and Virus Strains | | |
| **BL21-CodonPlus (DE3)-RIL Competent Cells, codon optimized protein expression strain,** $Cam^r$ (50 µg/ml) | Agilent Technologies | Cat#230245 |
| MP304: BL21-CodonPlus (DE3)-RIL+pMP35, *F341_rpb* protein expression strain, $Kan^r$ (100 µg/ml), $Cam^r$ (50 µg/ml) | This study | N/A |
| *Campylobacter jejuni* NCTC12658: Phage propagation strain | National collection of type cultures | N/A |
| *Campylobacter jejuni* NCTC12658Δ*motA: non-motile NCTC12658, paralyzed flagella, Kan*$^r$ (50 µg/ml) | Baldvinsson et al., 2014 | N/A |
| *Campylobacter jejuni* NCTC12658Δ*05515-05525 (*NCTC12658Δ*LOS), LOS mutant, Kan*$^r$ (50 µg/ml) | This study | N/A |
| LJ1: Motile phage F341 resistant *C. jejuni* NCTC12658 variant | This study | N/A |
| LJ4: Motile phage F341 resistant *C. jejuni* NCTC12658 variant | This study | N/A |
| LJ6: Motile phage F341 resistant *C. jejuni* NCTC12658 variant | This study | N/A |
| LJ8: Motile phage F341 resistant *C. jejuni* NCTC12658 variant | This study | N/A |
| LJ9: Motile phage F341 resistant *C. jejuni* NCTC12658 variant | This study | N/A |
| LJ10: Motile phage F341 resistant *C. jejuni* NCTC12658 variant | This study | N/A |
| LJ11: Motile phage F341 resistant *C. jejuni* NCTC12658 variant | This study | N/A |
| LJ13: Motile phage F341 resistant *C. jejuni* NCTC12658 variant | This study | N/A |
| *Campylobacter* phage F341, isolated in Denmark in 2004 from broiler intestine, *Fletchervirus,* Genbank: OQ864999 | This paper, Hansen et al., 2007 | N/A |
| Chemicals, Peptides, and Recombinant Proteins | | |
| Blood agar Base no. 2 | Oxoid | Cat#CM0271 |
| Brain Heart Infusion broth | Oxoid | Cat#CM1135 |
| NZCYM Medium | Sigma | Cat#N3643 |
| DNase I (1 U/µl) | ThermoFisher Scientific | Cat#EN521 |
| RNase A (10 mg/ml) | ThermoFisher Scientific | Cat#EN0531 |
| Proteinase K (20 mg/ml) | ThermoFisher Scientific | Cat#AM2546 |
| Glycogen | ThermoFisher Scientific | Cat#R0561 |
| Ammonium acetate | Sigma | Cat#A1542 |
| SmaI | New England Biolabs | Cat#R0141S |
| KpnI | New England Biolabs | Cat#R0142 |
| Critical Commercial Assays | | |
| MagAttract HMW DNA kit | Qiagen | Cat#67563 |
| Maxwell 16 Tissue DNA extraction kit | Promega | Cat#AS1030 |
| Oligonucleotides | | |

| | | |
|---|---|---|
| Primer: LOS_F_up: CATGCTCCTCTAGACTCGAGCAATACACTCAAAAAG AACGCGAT | This paper | N/A |
| Primer: LOS_R_up: AATGGTTCGCTGGGTTTATCTACGATGATAATAGCAC TTATTTGCTTTAGA | This paper | N/A |
| Primer: Kan+LOS_F: TAAGTGCTATTATCATCGTAGATAAACCCAGCGAACC ATTTGAG | This paper | N/A |
| Primer: Kan+LOS_R: TCACCTATGATACAACAATGCTAAAACAATTCATCCA GTAAAATATAATATTTTATTTTCTCC | This paper | N/A |
| Primer: LOS_F_down: TACTGGATGAATTGTTTTAGCATTGTTGTATCATAGG TGAAGCTTGAT | This paper | N/A |
| Primer: LOS_R_down: CGTTGGGAGCTCTCCGGATCCCATCCATTCTAGGTAC ACATTCTT | This paper | N/A |
| Primer: LOS_control_F: CACCGCAAAATCATCAATACAAAT | This paper | N/A |
| Primer: LOS_control_F: AACGACTATGATGCTTGAAATAACTT | This paper | N/A |
| Recombinant DNA | | |
| Plasmid: pET28a+: Protein expression vector: N-His, N-Thrombin, C-His, $Kan^r$ (100 µg/ml) | Novagen | N/A |
| Plasmid: pMP35: *F341_rbp* expression plasmid, pET28a+::*F341_rpb+chaperone*, $Kan^r$ (100 µg/ml) | This paper | N/A |
| Plasmid: pGEM-7Zf: vector for homologous recombination, $Amp^r$ (100 µg/ml) | This paper | N/A |
| Plasmid: pBCα3: vector containing *AphA-3* cassette giving kanamycin resistance, $Kan^r$ (50 µg/ml) | Bijlsma et al., 1999 | N/A |
| Plasmid: pMP810: pGEM-7Zf+::*05515-05525 deletion* construct, $Amp^r$ (100 µg/ml), $Kan^r$ (100 µg/ml) | This paper | N/A |
| Software and Algorithms | | |
| CLC genomics main workbench 20 | Qiagen | N/A |
| CPT Galaxy and WebApollo platforms | Afgan et al., 2018 | https://cpt.tamu.edu/galaxy-pub |
| InterPro | Mitchell et al., 2019 | https://www.ebi.ac.uk/interpro/ |
| HHpred | Zimmermann et al., 2018 | https://toolkit.tuebingen.mpg.de/tools/hhpred |
| Dali | Holm, 2022 | http://ekhidna2.biocenter.helsinki.fi/dali/ |
| BLAST | Johnson et al., 2008 | https://blast.ncbi.nlm.nih.gov/Blast.cgi |
| Colab AlphaFold2 | Mirdita et al., 2022 | https://colab.research.google.com/github/sokrypton/ColabFold/blob/main/AlphaFold2.ipynb |
| Pymol | Schrodinger, 2015 | N/A |
| Other | | |
| His GraviTrap | GE Healthcare Life Sciences | Cat#GE11-0033-99 |
| Amicon Ultra-15 Centrifugal filter units, 10 kDa | Merck | Cat#UFC901008 |

**Table S2. Single nucleotide polymorphisms (SNPs) observed by basic variant detection analysis of motile F341 resistant NCTC12658 variants.** Related to Table 1.

| | | | | | SNP, SNP frequency and resulting effect | | | | | | |
|---|---|---|---|---|---|---|---|---|---|---|---|
| **SNP position** | **Gene and size (locus)** | **Product (homolog in *C. jejuni* NCTC11168)** | **SNP description[a)]** | **NCTC12658 reference seq.[b)] (SNP location in gene)** | **LJ1** | **LJ6** | **LJ8** | **LJ9** | **LJ10** | **LJ11** | **LJ13** |
| 47.223 | *K5A08_00175* 3.747 bp | Eco57I restriction-modification methylase domain-containing protein (Cj0031, type IIG restriction-modification enzyme) | PolyG tract variation | 9 G's – on (2.573 bp) | 10 G's (79%) - off | 10 G's (85%) - off | 10 G's (85%) - off | 10 G's (90%) - off | 10 G's (92%) - off | 10 G's (92%) - off | 10 G's (60%) - off |
| 64.022 | *K5A08_00240* 765 bp | Bacteriohemerythrin (Cj0045, putative iron binding protein) | PolyG tract variation | 10 G's – on[c)] (717 bp) | 11 G's (85%), truncation: 241 aa vs 254 aa | 11 G's (53%), truncation: 241 aa vs 254 aa | 12 G's (61%), truncation: 240 aa vs 254 aa | | 11 G's (84%), truncation: 241 aa vs 254 aa | 11 G's (58%), truncation: 241 aa vs 254 aa | |
| 371.802 | *K5A08_01925* 546 bp | DUF1882 domain-containing protein (Cj0403, hypothetical protein) | SNV | (299 bp) | T to C (100%), F to S aa change | T to C (100%), F to S aa change | T to C (98%), F to S aa change | T to C (100%), F to S aa change | T to C (100%), F to S aa change | T to C (100%), F to S aa change | T to C (100%), F to S aa change |
| 530.435 | | Intergenic region | PolyG tract variation | 10 G's | 11 G's (59%) | 11 G's (81%) | | 11 G's (71%) | 11 G's (65%) | 11 G's (85%) | 11 G's (75%) |
| 1.078.122 | *wlaN* 911 bp (LOS) | Beta-1,3-galactosyltransferase (WlaN, beta-1,3 galactosyltransferase) | PolyG tract variation | 7 G's – off (337 bp) | | 9 G's (100%) – off | 9 G's (100%) – off | 9 G's (100%) – off | 9 G's (100%) - off | 9 G's (100%) – off | 9 G's (92%) – off |
| 1.230.874 | *K5A08_06310* 1.308 bp (OLG) | DUF4910 domain-containing protein (Cj1295, di-O-methylglyceroyl modification of Pse) | PolyG tract variation | 9 G's – on (144 bp) | | | 10 G's (82%) – off | | | | |
| 1.253.815 | (OLG) | Intergenic region | SNV | | | | | T to A (40%) | | | |

| | | | | | | | | | | | |
|---|---|---|---|---|---|---|---|---|---|---|---|
| 1.368.107 | *K5A08_06960* 639 bp (CPS) | Hypothetical protein | PolyG tract variation | 8 G's – off (127 bp) | | | | | | | 9 G's (100%) – on |
| 1.490.624 | *K5A08_07545* 1.956 bp | Methyl-accepting chemotaxis protein (Cj1564, putative methyl-accepting chemotaxis signal transduction protein) | SNV | (1.593 bp) | T to C (100%), no aa change | T to C (100%), no aa change | T to C (98%), no aa change | T to C (100%), no aa change | T to C (100%), no aa change | T to C (98%), no aa change | T to C (100%), no aa change |
| 1.594.549 | *K5A08_08145* 3.431 bp | Autotransporter outer membrane beta-barrel domain-containing protein (Cj0628, putative lipoprotein) | PolyG tract variation | 10 G's – off (497 bp) | | | | | | | 11 G's (38%) – on |

**Table S3. Gaps detected in genomes of phage F341 resistant NCTC12658 variants following basic variant detection (BVD) analysis (CLC Genomics Workbench 21.0.3, default settings).** Related to Table 1.

| **Gap position (locus)** | **Region or genes affected** | **Product (homolog in *C. jejuni* NCTC11168)** | **Description of gap and region in the *de novo* assembly** |
|---|---|---|---|
| **LJ1** | | | |
| 1.077.500..1.077.690 (LOS) | *K5A08_05530* (1.076.390..1.077.559) *wlaN* (1.077.548..1.078.458) | Glycosyltransferase family 2 protein (Cj1138) Beta-1,3-galactosyltransferase WlaN (WlaN) | BVD analysis: part at the end of the genes have no read coverage, *de novo* assembly: contig break |
| **LJ6** | | | |
| 1.077.608..1.077.687 (LOS) | *wlaN* (1.077.548..1.078.458) | Beta-1,3-galactosyltransferase WlaN (WlaN) | BVD analysis: minor part in the end of the gene has no read coverage, *de novo* assembly: contig break |
| 1.467.869..1.467.896 | *K5A08_07435* (1.467.547..1.468.542) | ABC transporter ATP-binding protein (Cj1538c) | BVD analysis: minor part in the middle of the gene has no read coverage, *de novo* assembly: contig break |
| **LJ8** | | | |
| 192.963..192.991 | *K5A08_00985* (192.427..193.548) | Hypothetical protein (Cj0199c) | BVD analysis: minor part in the middle of the gene has no read coverage, *de novo* assembly: contig break |
| 297.762..298.136 | *K5A08_01530* (297.431..298.627) | Hypothetical protein. Note: HHpred analysis: Triphosphate tunnel metalloennzyme, 1-368 aa (protein 398 aa), probability: 100, E-value: 8.5e-30, target length: 435. (Cj0323) | BVD analysis: middle of the gene has no read coverage, *de novo* assembly: contig break |
| 580.846..580.864 | *K5A08_03040* (580.082..581.312) | DUF2920 family protein (Cj0617 and Cj0618) | BVD analysis: minor part close to the middle of the gene has no read coverage, *de novo* assembly: contig break |
| 590.634..590.776 | Intergenic region | | BVD analysis: no read coverage upstream a hypothetical protein and putative lipoprotein both also found in another position on the genome, *de novo* assembly: contig break |
| **LJ9** | | | |
| 93.865..93.966 | Intergenic region | | BVD analysis: minor part has no read coverage, *de novo* assembly: contig break |
| 144.107..144.262 | Intergenic region *K5A08_00715* (144.213..146.231) | Methyl-accepting chemotaxis protein | BVD analysis: no read coverage across the intergenic region upstream and the first 50 bases of the gene, *de novo* assembly: contig break |
| 297.816..297.883 297.928..298.049 | *K5A08_01530* (297.431..298.627) | Hypothetical protein. Note: HHpred analysis: Triphosphate tunnel metalloennzyme, 1-368 aa (protein 398 aa), probability: 100, E-value: 8.5e-30, target length: 435. (Cj0323) | BVD analysis: two minor parts in the middle has no read coverage, *de novo* assembly: contig break |
| 380.813..380.882 | *K5A08_01965* (379.136..381.322) | Dynamin family protein (Cj0411) | BVD analysis: minor part end of the gene has no read coverage, *de novo* assembly: contig break |
| 434.275..434.452 | *nssR* (434.072...434.668) | Nitrosative stress-sensitive transcriptional regulator NssR (NssR) | BVD analysis: no read coverage in the middle of the gene, *de novo* assembly: contig break |

| | | | |
|---|---|---|---|
| 580.975..581.000 | *K5A08_03040* (580.082..581.312) | DUF2920 family protein (Cj0617 and Cj0618) | BVD analysis: minor part middle of the gene has no read coverage. *de novo* assembly: contig break |
| 590.634..590.752 | Intergenic region | | BVD analysis: no read coverage upstream *K5A08_03090* encoding an autotransporter outer membrane beta-barrel domain-containing protein also found in another position on the genome (*K5A08_08145)*, *de novo* assembly: contig break |
| 814.579..814.603 814.691..814.788 | Intergenic region | | BVD analysis: two minor parts has no read coverage, *de novo* assembly: contig break |
| 925.125..925.137 | Intergenic region | | BVD analysis: minor part has no read coverage, *de novo* assembly: contig break |
| 996.561..996.573 | *K5A08_05120* (994.856..996.829) | LTA synthase family protein (Cj1055c) | BVD analysis: very minor gap with no read coverage, *de novo* assembly: contig break |
| 1.032.947..1.032.954 | *Int* (1.031.983..1.033.308) | Apolipoprotein N-acyltransferase (Int) | BVD analysis: very minor gap with no read coverage, *de novo* assembly: contig break |
| 1.077.562..1.077.605 (LOS) | *wlaN* (1.077.548..1.078.458) | Beta-1,3-galactosyltransferase WlaN (WlaN) | BVD analysis: minor part at the end of the gene has no read coverage, *de novo* assembly: contig break |
| 1.363.624..1.363.643 (CPS) | *K5A08_06930* (1.361.818..1.363.736) | Sugar transferase | BVD analysis: minor part in the beginning of the gene has no read coverage, *de novo* assembly: contig break |
| 1.467.717..1.467.842 | *K5A08_07435* (1.467.547..1.468.542) | ABC transporter ATP-binding protein (Cj1538c) | BVD analysis: part at the end of the gene has no read coverage, *de novo* assembly: contig break |
| 1.507. 499..1.507.593 | *K5A08_07625* (1.506.938..1.507.603) | ATP-binding cassette domain-containing protein (Cj1580c) | BVD analysis: part at the beginning of the gene has no read coverage, *de novo* assembly: contig break |
| **LJ10** | | | |
| 1.074.429..1.076.390 (LOS) | *K5A08_05520* (1.074.181-1.075.353) *K5A08_05525* (1.075.337..1.076.332) | Glycosyltransferase family 2 protein (Cj1136) Capsular polysaccharide synthesis protein (Cj1137) | BVD analysis: part has no read coverage (most of *K5A08_05520* and all of *K5A08_05525* have no read coverage), *de novo* assembly: contig break, unclear if genes are missing |
| **LJ11** | | | |
| 814.710..814.816, 814.869..814.970 | Intergenic region | | BVD: part has no read coverage, *de novo* assembly: contig break |
| **LJ13** | | | |
| 48.905..48.990 | *K5A08_00180* (48.397..50.214) | Pentapeptide repeat-containing protein | BVD analysis: part has no read coverage, *de novo* assembly: contig break |
| 144.001..144.095 | Intergenic region | | BVD analysis: part has no read coverage, *de novo* assembly: contig break |
| 1.074.820..1.076.414 (LOS) | *K5A08_05520* (1.074.181-1.075.353) *K5A08_05525* (1.075.337..1.076.332) *K5A08_05530* (1.076.390..1.077.559) | Glycosyltransferase family 2 protein (Cj1136) Capsular polysaccharide synthesis protein (Cj1137) Glycosyltransferase family 2 protein (Cj1138) | BVD analysis: most of *K5A08_05520*, all of *K5A08_05525* and first 25 bases of *K5A08_05530* have no read coverage, *de novo* assembly: The *K5A08_05520* and *K5A08_05525* genes are no longer present in the variant |

**Table S4. Genes unique to phage F341.** Related to Table 2.

| Gene | Size (bp) | Product | Size (aa) | BLASTN suite | BLASTP suite | Domains (InterPro) | Domains (HHpred) | AlphaFold2 prediction and Dali search |
|---|---|---|---|---|---|---|---|---|
| *F341_109* | 516 | Putative adaptor protein | 171 | No significant similarity | Putative carbohydrate binding protein F336_112, *Campylobacter* phage F336, Query cover: 100%, E-value: 5e-57, Per. Ident.: 55.8% | Carbohydrate-binding module superfamily 5/12, 54-97 aa | Short-tailed cyanophage tailspike receptor-binding domain, 45-98 aa, Probability: 93.2%, E-value: 0.2, Score: 47.2% | Baseplate wedge protein, adaptor protein |
| *F341_128* | 1.470 | Putative baseplate wedge protein | 489 | *Campylobacter* phage PC14, *PC14_00096* hypothetical protein, Query cover: 79%, E-value: 2e-149, Per. Ident.: 75,4% | Hypothetical protein F207_121, *Campylobacter* phage F207, Query cover: 96%, E-value: 0.0, Per. Ident.: 60.6% | No domains detected | Baseplate wedge protein gp7, Enterobacteria phage T4, 7-195 aa, Probability: 78.8%, E-value: 11, Score: 44.3%, | Poor models, no structural evidence to support HHpred proposed function |
| *F341_135* | 309 | Putative fibritin | 102 | No significant similarity | Phage tail protein, *Clostridioides difficile*, Query cover: 83%, E-value: 1e-04, Percent Identity: 38.2%, | No domains detected | Fibritin, Bacteriophage T4, 10-101 aa, Probability: 54.9%, E-value: 55, Score: 28.3% | Fibritin |
